# The A53T mutation in α-synuclein enhances pro-inflammatory activation in human microglia

**DOI:** 10.1101/2023.08.29.555300

**Authors:** Marine Krzisch, Bingbing Yuan, Wenyu Chen, Tatsuya Osaki, Dongdong Fu, Carrie Garrett-Engele, Devon Svoboda, Kristin Andrykovich, Mriganka Sur, Rudolf Jaenisch

## Abstract

Parkinson’s disease (PD) is characterized by the aggregation of α-synuclein into Lewy bodies and Lewy neurites in the brain. Microglia-driven neuroinflammation may contribute to neuronal death in PD, however the exact role of microglia remains unclear and has been understudied. The A53T mutation in the gene coding for α-synuclein has been linked to early-onset PD, and exposure to A53T-mutant human α-synuclein increases the potential for inflammation of murine microglia. To date, its effect has not been studied in human microglia. Here, we used 2-dimensional cultures of human iPSC-derived microglia and transplantation of these cells into the mouse brain to assess the effects of the A53T mutation on human microglia. We found that A53T-mutant human microglia had an intrinsically increased propensity towards pro-inflammatory activation upon inflammatory stimulus. Additionally, A53T mutant microglia showed a strong decrease in catalase expression in non-inflammatory conditions, and increased oxidative stress. Our results indicate that A53T mutant human microglia display cell-autonomous phenotypes that may worsen neuronal damage in early-onset PD.

## Introduction

Parkinson’s disease (PD) is the second most common neurodegenerative disease, after Alzheimer’s disease. It is characterized by the aggregation of α-synuclein into Lewy bodies and Lewy neurites in the brain. Microgliosis is an early and sustained response in PD [1, 2], and microglia-driven neuroinflammation may contribute to neuronal death in PD. However, the exact role of microglia remains unclear and has been understudied.

The A53T mutation is a missense point mutation in the gene encoding α-synuclein, *SNCA*, and has been linked to autosomal dominant, early-onset PD [3]. A53T α-synuclein fibrillizes faster than wild-type (WT) alpha synuclein [4], and exposure of mouse microglia to human A53T-mutant α-synuclein promotes their inflammatory cascades more strongly than exposure to human WT α-synuclein [5]. Furthermore, microglia develop a strong reactive state with an excessive production of reactive oxygen species (ROS) and pro-inflammatory cytokines in a mouse model with selective α-synuclein accumulation in microglia [6]. However, to our knowledge, the effect of the A53T α-synuclein mutation in human microglia has not yet been studied.

Mouse models fail to fully recapitulate the pathogenesis of PD [7], and, so far, have failed to yield a cure to this disease. Furthermore, a recent study showed that, although mouse and human microglia are largely similar, they age differently [8], and human microglia express genes relevant to human neurodegenerative disease that are not expressed by other mammals. This highlights the need to study human-relevant pathways using human microglia [9].

Culture conditions fail to fully recapitulate the physiological characteristics of human microglia [10, 11]. Myeloid precursors (MPs) derived from human pluripotent stem cells (hPSC) and transplanted into the brain of human CSF1 knock-in immune-deprived mouse neonates colonize the mouse brain and yield microglia that retain their human identity, and more closely resemble *ex vivo* human microglia than *in vitro* 2D cultures [12-15]. Hence, transplantation of human MPs carrying PD-related mutations in the mouse brain may yield new, more physiologically relevant insights on the dysfunction of microglia in PD.

Here, we used A53T-mutant human pluripotent stem cells (hPSC) and isogenic controls to characterize the effects of the A53T mutation on human microglia. After screening for potential phenotypes using bulk RNA sequencing of cultured human microglia, we generated mice carrying A53T-mutant (PD) or isogenic control GFP-labeled human microglia in their brain by transplanting hPSC-derived MPs in the brain of human CSF1 knock-in immune-deprived mouse neonates and letting them differentiate into microglia inside the mouse brain, in order to study their morphological and functional alterations in a more physiologically relevant context. Our results indicate that PD microglia had decreased levels of expression of catalase, a ROS scavenger, in non-inflammatory conditions, and increased levels of oxidative stress. Furthermore, they displayed higher levels of pro-inflammatory activation when placed in pro-inflammatory conditions. This may contribute to neuronal damage in early-onset PD.

## Material and methods

### Experimental Animals

Rag2/IL2rg double knock-out mice carrying a transgene for the human allele of CSF1 (C;129S4-Rag2tm1.1Flv Csf1tm1(CSF1) Flv Il2rgtm1.1Flv/J) were obtained from the Jackson Laboratory. Mice were kept in group housing in under standard barrier conditions. Food and water were available *ad libitum*. All experiments were performed in accordance with the Department of Comparative Medicine and Massachusetts Institute of Technology standards.

### hPSC culture and labeling

hPSC lines carrying the A53T mutation in α-synuclein and control cell lines used in this study were previously generated and characterized by our group and are listed in Table 1 [16]. All cell lines were targeted with the Green Fluorescent Protein (GFP) at the AAVS1 locus with the plasmid available on Addgene (#22212) using the Zinc-Finger approach as previously described [17]. hPSCs were then switched to feeder-free conditions. hPSCs were regularly tested for mycoplasma contamination. To prevent accidental inversion of the cell lines, genotyping for the A53T mutation was regularly performed at the beginning of each differentiation.

### In vitro differentiation of microglia

*In vitro* differentiation of microglia was performed using a modified version of a published protocol [18]. Briefly, hPSCs were cultured in mTESR1 and differentiated into hematopoietic progenitor cells (HPCs) using the Stemdiff™ hematopoietic kit. On day 10-12 of differentiation, HPCs were plated at 10^5^ cells per well on Matrigel coated 6-well plates in microglial differentiation medium (basal medium supplemented with 200 ng/mL IL-34, 100 ng/mL TGFβ-1, and 50 ng/mL M-CSF). 1 mL of microglial differentiation medium per well was added every other day for 25 days. On day 12 of differentiation, 5 mL of medium were removed from each well and centrifuged. The cell pellet was resuspended in fresh microglial differentiation medium (1 mL per well) and plated back. On day 25, 5 mL of medium were removed from each well and centrifuged. The cell pellet was resuspended in fresh microglial maturation medium (1 mL per well) and plated back. Microglial maturation medium consisted of basal medium supplemented with 100 ng/mL CD200, 100 ng/mL CX3CL1, 200 ng/mL IL-34, 100 ng/mL TGFβ-1, and 50 ng/mL M-CSF. On day 27, 1 mL of microglial maturation medium was added to each well. On day 29, microglia were harvested for characterization and experimentation.

### MP differentiation and transplantation

hPSCs were differentiated into MPs using a previously published protocol [19]. MPs were released into suspension after two weeks in hematopoietic medium and these cells were collected during each medium change within 1.5 month of culture in hematopoietic medium for transplantation into neonatal mice. Cultured MPs were resuspended in phosphate-buffered saline without calcium and magnesium prior to injection at a concentration of 10^5^ cells/µL. Postnatal day 0 (P0) to P3 mouse pups of either sex were manually injected with 4.10^5^ MPs in the lateral ventricles (1 anterior and 1 posterior injection site per brain hemisphere) using glass micropipettes.

### Transplanted cell extraction for RNA sequencing

Transplanted cells were extracted from the mouse brains at 2 months post-transplantation as previously described [13]. The cells were stained with DAPI prior to Fluorescence-activated cell sorting (FACS) sorting for GFP and DAPI, and only DAPI-negative GFP-positive cells were retained for analysis.

### RNA sequencing and analysis

RNA from human microglia differentiated *in vitro* and human microglia extracted from human-mouse chimeras was extracted using the RNeasy Plus Micro Kit (Qiagen). The RNA from 3 independent *in vitro* microglia differentiations per cell line were pooled at equal weight to obtain six samples, each corresponding to a cell line. Similarly, the RNA from transplanted cells extracted from 2 to 3 chimeras per cell line were pooled at equal weight. RNA libraries were prepared using NEBNext® Single Cell/Low Input RNA Library Prep Kit for Illumina® (New England Biolabs) and sequenced using the Illumina NOVAseqSP sequencer.

The Illumina paired-end reads were mapped with STAR [20] (version 2.7.1a), using index files created with the gene annotation from Ensembl human hg38 version 108 and mouse mm10 version 102 with the ‘--sjdbOverhang 50’ option. Genes counts were obtained using the featureCounts [21] function from Subread package with the unstranded option. Gene expression levels were normalized by library size. Differential expression analysis, based on the negative binomial distribution, was done with DESeq2 [22] with a paired design and local fitting of dispersions to the mean intensity. To accurately estimate the logarithmic fold change (LFC) for genes with low expression levels or with high variation, an adaptive normally distributed prior was applied for LFC shrinkage. Genes with FDR-adjusted p-value<0.05 and at least two-folds difference were considered to be differentially expressed. Raw data along with gene expression levels were deposited to NCBI Gene Expression Omnibus (GSE241437). We performed gene list enrichment analysis on the differentially expressed genes using ToppFun from the ToppGene suite [23, 24]. For Gene Set Enrichment Analysis (GSEA) [25] Preranked, we ranked the genes by the statistical test statistic, including only protein coding genes. The analysis was done on the Hallmark gene sets collection (h.all.v2023.1.Hs.symbols.gmt) for comparisons between *in vitro* differentiated and transplanted microglia, and on the Hallmark and curated (c2.all.v2023.1.Hs.symbols.gmt) gene set collections, both from MSigDB.

### Lipopolysaccharide (LPS) injections

At 2 months post-transplantation, transplanted mice were given a single intraperitoneal injection of LPS (Sigma Aldrich L2630) dissolved in sterile saline (5mg/kg). Animals were euthanized 24 hours after the injection and brains were collected for immunofluorescence analyses.

### Immunofluorescence staining

Cells in culture were fixed using 4% paraformaldehyde. Mice were perfused with 4% paraformaldehyde through the transcardial route and brains were harvested. 100 µm-thick (for percentages of positive cells) and 20 µm-thick (for signal intensity measurements) brain sagittal cryosections were generated after cryopreservation in 30% sucrose. Immunostainings were performed using antibodies listed in Tables 2 and 3.

### Confocal imaging and analyses

Images were acquired using a Zeiss LSM710 confocal microscope (tile scan image) and a Zeiss LSM700 confocal microscope (other images). To assess microglia morphology and the percentage of marker-positive cells, Confocal microscopy z-stacks of control and PD microglia were acquired with a 20x dry objective (zoom 0.6), z-step of 2 µm. Maximum intensity projections were generated using the Fiji software (ref) and analyzed. To assess the intensity of fluorescence signal, confocal microscopy z-stacks were acquired using a 20x dry objective (zoom 1), z-step of 0.95 µm. 3D images were analyzed using the Imaris software (9.9.1). Microglial cells were surfaced using the Surfaces function, and the integrated density of the fluorescence signal was calculated using the following formula: mean intensity of signal x number of voxels.

### Statistical analyses of immunofluorescence experiments

Statistical analyses were performed using GraphPad Prism (v.9) (GraphPad software, San Diego, California, USA, www.graphpad.com). Error bars in the figures indicate standard error of the mean (SEM). For the analysis of microglial morphological categories, we performed a two-way repeated measures ANOVA test or a mixed-effects model followed by a Šídák correction for multiple comparisons. For other analyses, when the number of values per cell lines was inferior or equal to 6, an unpaired t-test was used. In other cases, distribution normality of each group of data was assessed with a Shapiro–Wilk test. When the distribution was not normal, the non-parametric Mann–Whitney test was applied. Otherwise, the equality of standard deviations of the groups was tested. If the groups had similar standard deviations, a standard unpaired -t-test was used. If not, an unpaired t-test with Welch’s correction was used.

## Results

### Human A53T-mutant microglia display gene signatures suggesting an altered state of activation

To generate hypotheses on the effects of the *SNCA* A53T mutation in human microglia, we first performed bulk RNA sequencing on PD and isogenic control microglia differentiated *in vitro*. 81 to 98% of *in vitro* differentiated cells were Iba1- and P2RY12-positive at day 29 of differentiation, indicating successful differentiation of all the cell lines into human microglia (Figure 1A, Table 2). We assessed the expression of key microglial genes, as defined in the literature [10, 13], in PD and control microglia. PD and control microglia expressed most key microglial genes at high levels, and no differences in the level of expression of these genes were found between PD and control microglia (Figure S1A). This confirmed the successful differentiation and similar differentiation stages of PD and control microglia, and indicated that the differences observed between PD and control microglia were unlikely to be due to different stages of differentiation.

**Figure 1.**
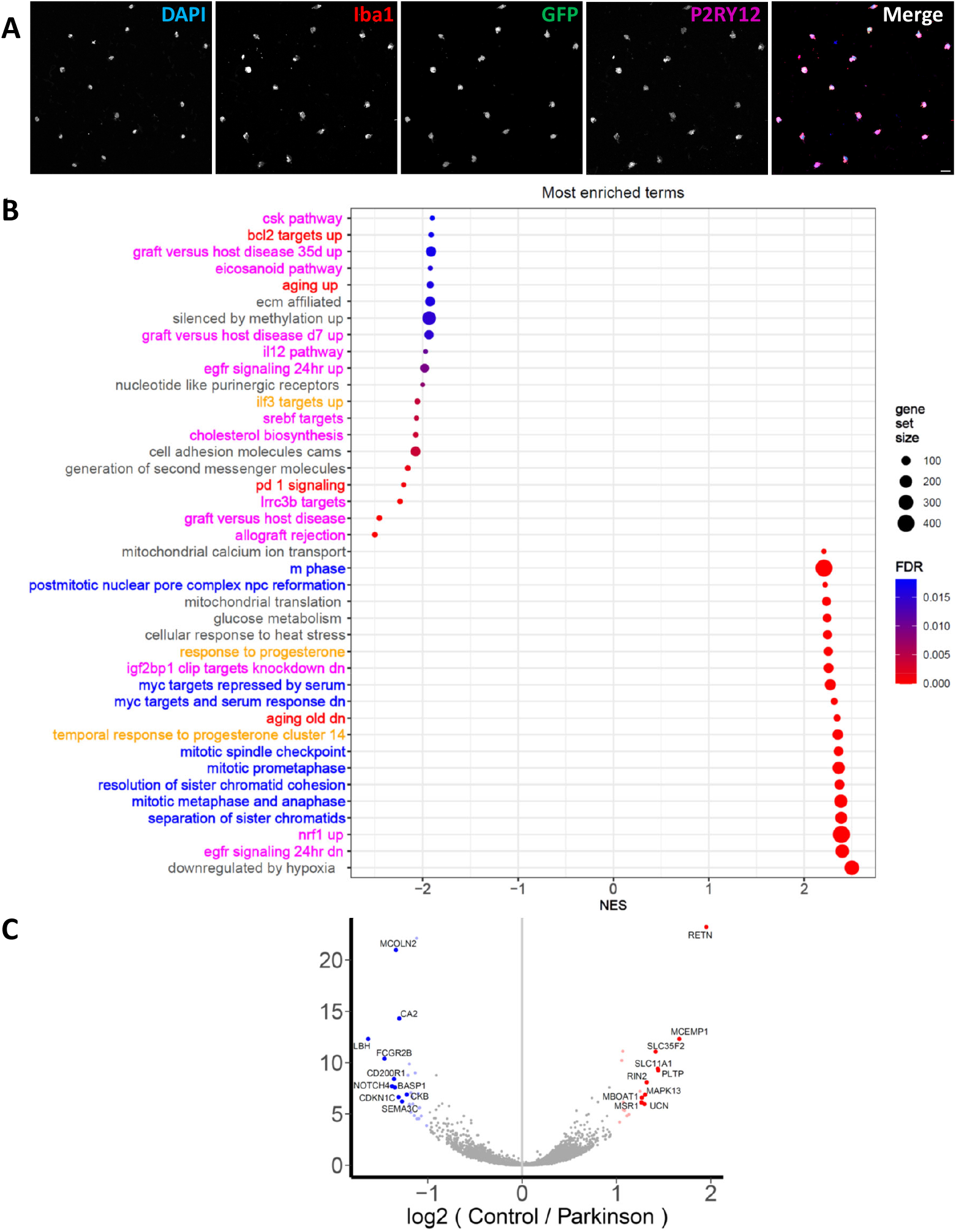
Bulk RNA sequencing analysis of hPSC-derived microglia differentiated *in vitro* shows differential expression of gene pathways linked to inflammation, immunomodulation, cell cycle and cellular senescence in PD microglia compared to isogenic control. **A**. Confocal maximum intensity projection of a 2-dimensional culture of GFP-labeled human pluripotent stem cell-derived microglia at 29 days *in vitro* (DIV), immunostained with DAPI (blue), Iba1 (red) and P2RY12 (magenta). Scale bar represents 20 µm. **B**. Upregulated and downregulated pathways in PD versus isogenic control *in vitro* differentiated microglia as assessed by GSEA. For clarity, only select pathways are shown. The exhaustive list of pathways can be found in Tables S1-S4. Normalized enrichment score (NES)>0 indicates upregulation of the pathway (lower panel), whereas NES<0 indicates downregulation of the pathway (upper panel) in PD microglia. The FDR threshold was set at 0.05. Gene pathways involved in immune response, inflammation and microglia activation are shown in magenta. Gene pathways involved in senescence and aging are shown in red. Gene pathways linked to immunomodulation are shown in orange. Gene pathways involved in cell cycle and cell proliferation are shown in blue. **C**. Volcano plot showing upregulated and downregulated genes in PD vs control *in vitro* differentiated microglia. The DE genes with adjusted p-value less than 0.05 and at least 2-fold differences are highlighted in red (upregulated genes) and blue (downregulated genes), and gene symbols for the top 10 genes are labeled. n=3 biological experiments for each cell line.

Assessment of the effect of the A53T mutation on *SNCA* expression levels has yielded conflicting results in the literature [26, 27]. In our study, the A53T mutation or its correction did not change the levels of expression of the *SNCA* gene in PD microglia compared to control (Figure S1B). Gene Set Enrichment Analysis (GSEA) showed an alteration of pathways involved in immune response, inflammation, microglia activation, and immune modulation in PD microglia (Figure 1B, Tables S1-S4). Additionally, pathways related to cell cycle were upregulated, and pathways related to cell death, senescence and aging were differentially expressed (Figure 1B, Tables S1-S4). The ten most downregulated genes in PD microglia included genes involved in immunomodulation: *FCGR2B, NOTCH4, CD200R1, SEMA3C* (Table 5), and the ten most upregulated genes in PD microglia included genes involved in immune response and associated to inflammatory diseases: *RETN, MCEMP1, SLC11A1, MAPK13, MSR1* (Table 6). Together, these results point to an altered state of activation of human PD microglia compared to isogenic control, increased proliferation and cellular senescence of PD microglia.

Primary microglia cultured in a dish lose their microglia-specific gene signatures and start expressing genes related to inflammation after six hours in culture. Transplanting human MPs into the mouse brain supports microglial differentiation and largely overcomes this limitation. Human PSC-derived microglia transplanted into the mouse brain assume a primary human microglia-like state, with key aspects of human microglial gene expression that cannot be recapitulated in culture [12-15].

To study the altered state of activation of PD microglia in this more physiologically relevant model, we transplanted PD hPSC-derived or isogenic control MPs into the mouse brain, allowing these cells to differentiate into microglia and mature in an *in vivo*-like environment (Figure 2A). At 2 months post-injection (PI), the transplanted cells had successfully populated the mouse brain (Figure 2B), and 93 to 100% of the cells were Iba1- and P2RY12-positive (Figure 2C, Table 7), indicating that they had successfully differentiated into human microglia.

**Figure 2.**
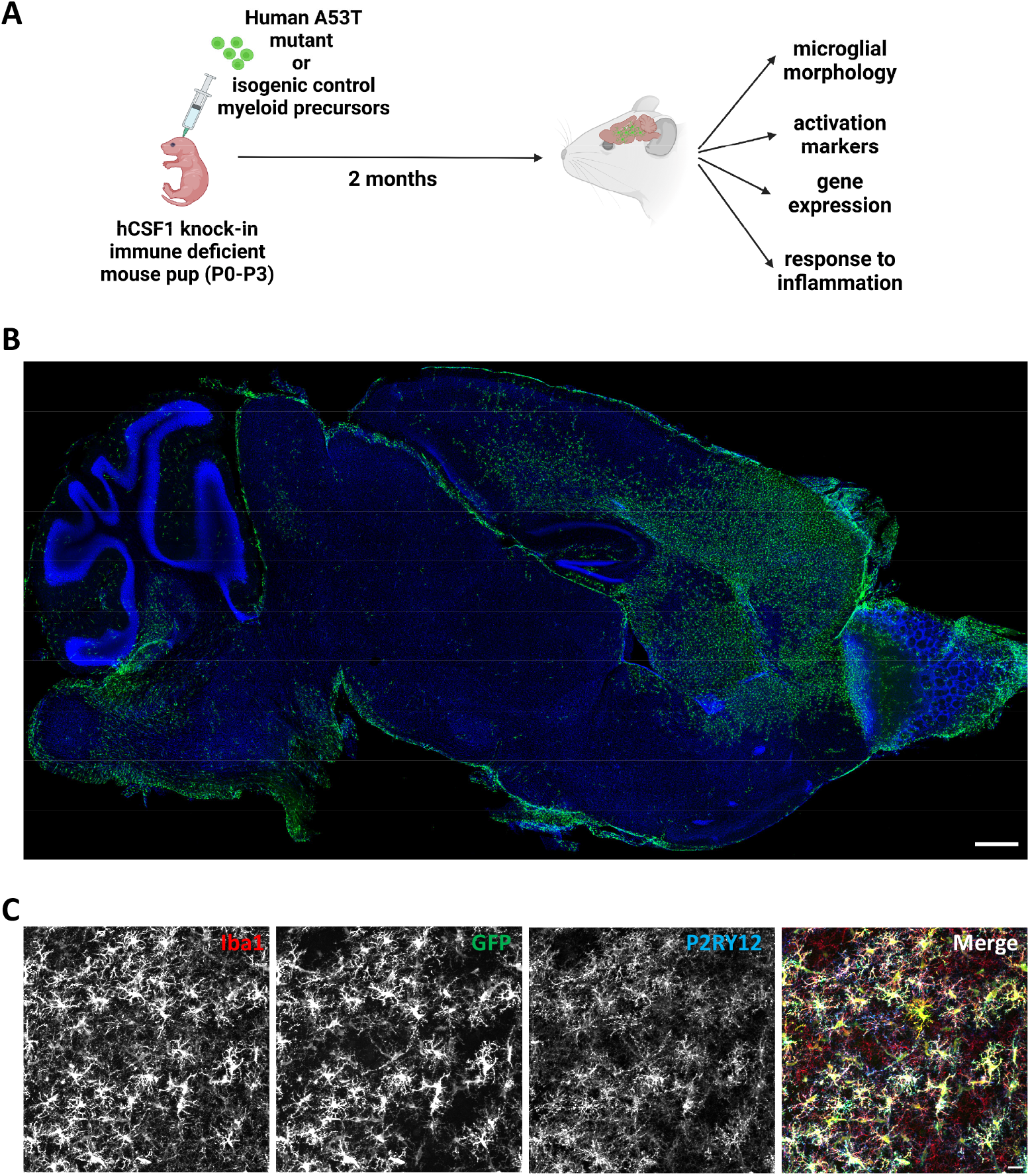
Transplanted hPSC-derived MPs have populated the mouse brain and differentiated into microglia at 2 months post-transplantation. **A**. Experimental design. For each isogenic pair of human pluripotent stem cell lines (hPSC), one cohort of immune-deficient neonatal mouse pups was injected with PD hPSC-derived myeloid precusors (MPs), and another cohort was injected with isogenic control hPSC-derived MPs. Mouse brains were analyzed at 2 months post-injection (PI). **B**. Tile scan confocal maximum intensity projection of a 100 µm-thick sagittal brain slice of a transplanted mouse at 2 months PI. Scale bar is 500 µm. Green: GFP-labeled hPSC-derived microglia, blue: DAPI. **C**. Confocal maximum intensity projection of transplanted hPSC-derived cells at 2 months PI, immunostained for microglia markers. Red: Iba1; green: GFP; blue: P2RY12. Scale bar is 20 µm.

At 2 months PI, PD and isogenic control microglia were extracted from the mouse brains and RNA sequencing was performed. Similar to *in vitro* differentiated PD microglia, transplanted PD microglia showed similar levels of expression of *SNCA* compared to control (Figure S1C). PD and control microglia expressed most key microglial genes at high and similar levels (Figure S1D). This confirmed the successful differentiation and similar differentiation stages of PD and control microglia, and indicated that the differences observed between PD and control microglia were unlikely to be due to different stages of differentiation.

To confirm that the transplantation of human microglia into the mouse brain led to a more physiologically relevant gene expression profile, we assessed the expression of a panel of key microglial genes previously assessed by our group [13]. The expression of the key microglial genes *CX3CR1, GPR34, P2RY12, P2RY13, SALL1, TGFBR1* and *TMEM119* was increased in transplanted microglia compared to *in vitro* differentiated microglia, in both control (Figure S2A) and PD cell lines (Figure S2B), indicating that transplanted microglia recapitulated human microglial gene expression profile to a higher degree than *in vitro* differentiated microglia. The expression of *ADGRG1* was, however, downregulated in transplanted microglia.

The expression of *C1QA*, a key microglial gene involved in microglia activation [28, 29], was slightly but significantly downregulated in both PD and control transplanted microglia compared to *in vitro* differentiated microglia. Additionally, the expression of genes coding for the pro-inflammatory cytokines IL1B and TNF was significantly downregulated, and the expression of the gene coding for IL10, an anti-inflammatory cytokine, was significantly upregulated in transplanted control microglia compared to *in vitro* differentiated control microglia. In PD transplanted microglia, *IL1B* and *TNF* also tended to be downregulated, although not significantly, and *IL10* was upregulated compared to PD *in vitro* differentiated microglia. Together, these results suggest lower levels of inflammatory response of control transplanted microglia compared to control *in vitro* differentiated microglia.

To confirm lower levels of inflammatory response in transplanted microglia compared to microglia differentiated *in vitro*, we assessed the differentially expressed pathways in *in vitro* differentiated microglia compared to transplanted microglia using gene list enrichment analysis (Figures S3, S4). Pathways related to immune response, inflammation and immune modulation were differentially expressed in *in vitro* differentiated microglia compared to transplanted microglia, and pathways linked to cell cycle were upregulated in PD *in vitro* differentiated microglia compared to PD transplanted microglia (Figures S3, S4). Together, these results indicate an alteration of the activation state in *in vitro* differentiated microglia compared to transplanted microglia.

To validate the hypotheses regarding the disease-relevant pathways of PD microglia as suggested by the results from *in vitro* differentiated microglia, we used GSEA to assess the differentially expressed pathways in transplanted PD microglia compared to transplanted control microglia. GSEA indicated an upregulation of pathways involved in translation, metabolism, immune response, inflammation, and microglia activation in PD microglia, and a downregulation of pathways involved in immune modulation, and DNA repair. In contrast to the results yielded by *in vitro* differentiation of microglia, pathways linked to cell cycle and cell proliferation were downregulated in PD transplanted microglia. Pathways related to cell death, cellular senescence and aging were differentially expressed in PD transplanted microglia compared to control (Figure 3A, Tables S5-S8).

**Figure 3.**
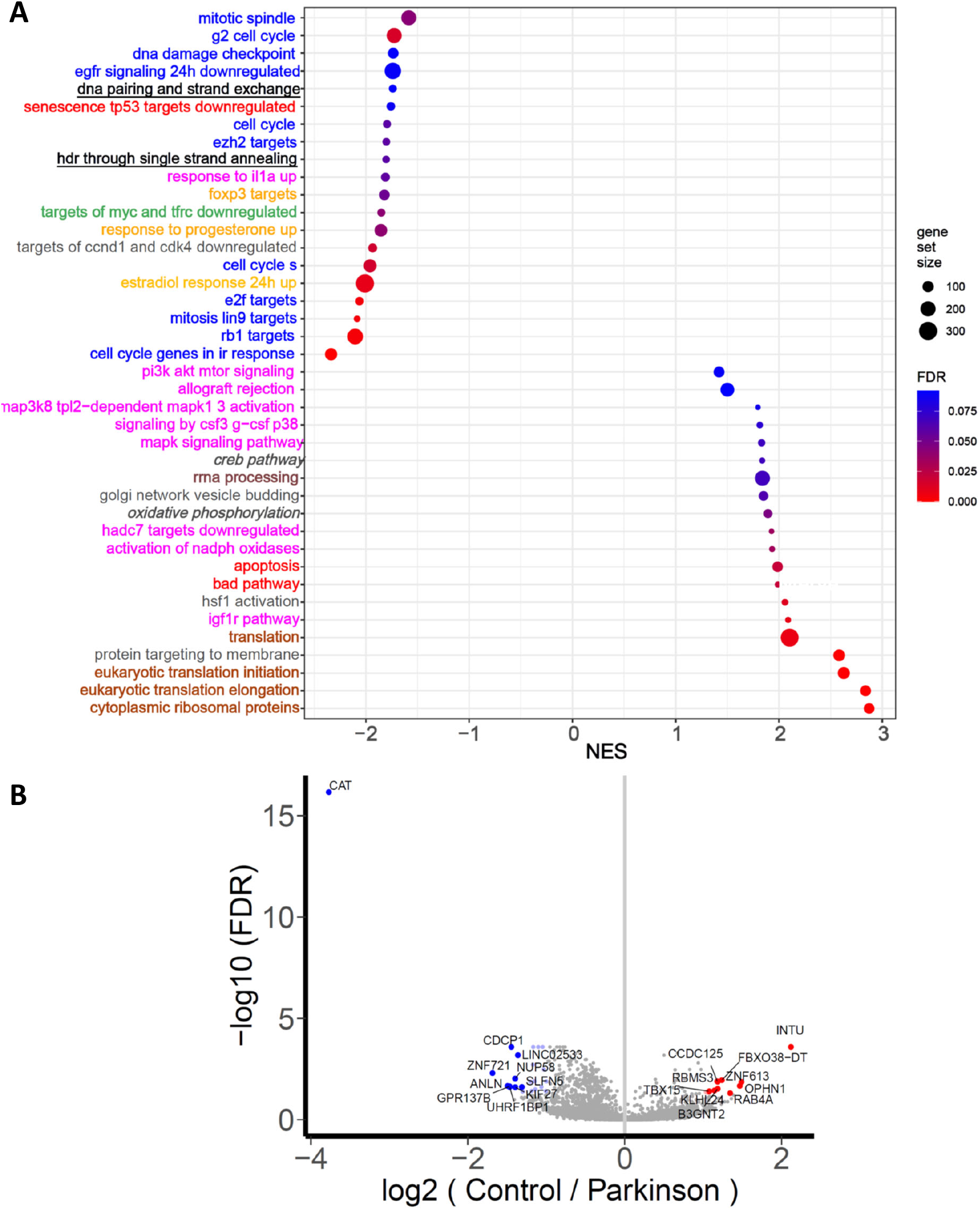
Bulk RNA sequencing analysis of hPSC-derived microglia transplanted into the brain of mice shows differential expression of gene pathways linked to inflammation, microglia activation, immune response, immunomodulation, protein translation, metabolism, cell cycle, DNA repair and cellular senescence in PD microglia compared to isogenic control. **A**. Upregulated and downregulated pathways in PD versus isogenic control transplanted microglia as assessed by GSEA. For clarity, only select pathways are shown. The exhaustive list of pathways can be found in Tables S5-S8. Normalized enrichment score (NES)>0 indicates upregulation of the pathway (lower panel), whereas NES<0 indicates downregulation of the pathway (upper panel) in PD transplanted microglia. The FDR threshold was set at 0.1. Gene pathways involved in immune response, inflammation and microglia activation are shown in magenta. Gene pathways involved in cell death, senescence and aging are shown in red. Gene pathways linked to immunomodulation are shown in orange. Gene pathways involved in cell cycle and cell proliferation are shown in blue. Gene pathways involved in translation are shown in brown. Gene pathways involved in DNA repair are shown in black, underlined. Gene pathways linked to metabolism are shown in black, italic. A gene pathway involved in cell cycle arrest is shown in green. **B**. Volcano plot showing upregulated and downregulated genes in PD vs control transplanted microglia. The DE genes with adjusted p-value less than 0.05 and at least 2-fold differences are highlighted in red (upregulated genes) and blue (downregulated genes), and gene symbols for the top 10 genes are labeled. n=2-3 transplanted brains per cell line.

The most differentially expressed gene in PD transplanted microglia was *CAT*, coding for the protein catalase, which was strongly downregulated (Figure 2B, Table 8). The *CAT* gene was also significantly downregulated in *in vitro* cultures, although to a lesser degree: log_2_FC(PD/control)=-0.2, p_adj_=0.04. Catalase mitigates oxidative stress by breaking down ROS [30]. The ten most significantly upregulated genes in PD microglia included *RBMS3*, which has been linked to motor complications in PD [31] (Table 9). Together, these results suggest that human PD microglia differentiated within the more physiological context of the mouse brain may display increased protein translation, metabolism, pro-inflammatory activation, oxidative stress and decreased cell division. Because of the differential expression of cell death, senescence and aging pathways, and the downregulation of pathways involved in DNA repair and cell cycle, PD transplanted microglia might also display increased cellular senescence.

### Human A53T-mutant microglia show increased pro-inflammatory activation in pro-inflammatory conditions

We then performed immunofluorescence analyses on transplanted mouse brain slices at 2 months PI. We focused on transplanted microglia located in the mouse striatum, as this region was consistently populated with human microglia in transplanted mice, and is involved in PD [32]. During activation, microglia undergo morphological changes. Quiescent microglia are characterized by a complex, ramified morphology. As microglia become activated, they undergo thickening and retraction of their branches, and acquire a bushy appearance, with fewer primary processes and poor ramifications. As activation progresses, they lose their ramifications and become ameboid [33]. CD68 labels lysosomes and is a widely used marker for microglia activation, as its expression increases in activated microglia [34]. Iba1 is the most commonly used marker of microglial activation [34]. p65 is a subunit of the NF-κB complex, which targets genes involved in inflammation development and progression [35]. The canonical NFκB p65/p50 pathway was shown to be activated in post-mortem PD human brains and the substantia nigra of animals undergoing dopaminergic neuron degeneration [36]. Additionally, NFκB canonical pathway activation leads to microglia pro-inflammatory activation and motor neuron death via inflammatory pathways [37].

We first separated microglia into different categories according to their morphology, using an approach similar to the approach undertaken by previous studies on microglia activation [38, 39] (Figure 4A). Quiescent microglia had a small cell body and a high number of ramifications. Bushy I microglia had enlarged cell bodies, with fewer processes. Bushy II microglia had enlarged cell bodies with less than ten primary processes. Ameboid microglia had round, enlarged cell bodies, with less than three primary processes. We then quantified percentages of CD68-positive and p65-positive PD and isogenic control transplanted microglia, and the staining intensity of Iba1 in PD and isogenic control transplanted microglia.

**Figure 4.**
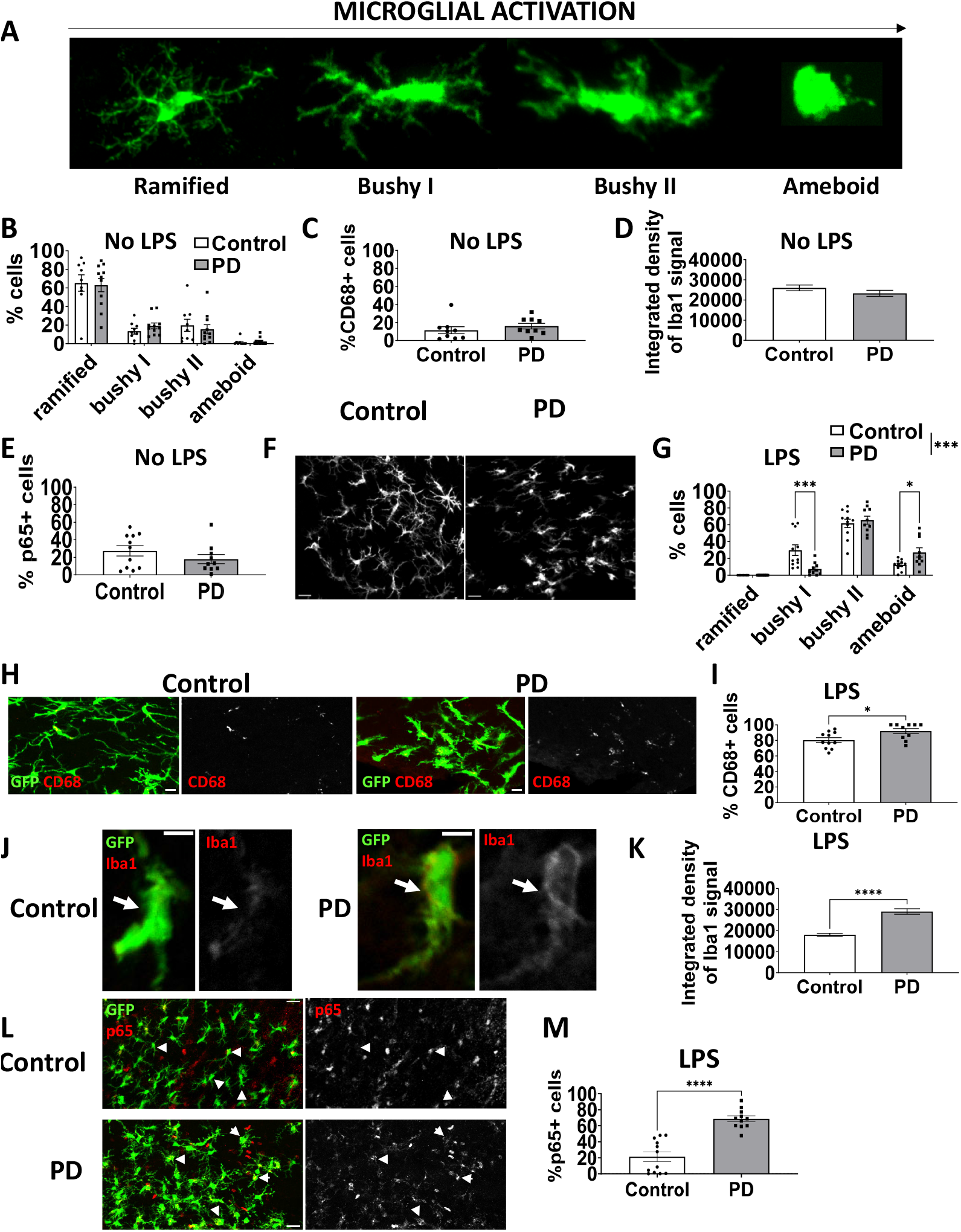
Transplanted striatal PD microglia display increased pro-inflammatory activation compared to isogenic control in pro-inflammatory conditions. **A**. Confocal maximum intensity projections illustrating the classification of transplanted human microglia into different morphological categories. Ramified microglia had a small cell body and a high number of ramifications. Bushy I microglia had enlarged cell bodies, with fewer processes. Bushy II microglia had enlarged cell bodies with less than ten primary processes. Ameboid microglia had round, enlarged cell bodies, with less than three primary processes. **B**. Percentages of ramified, bushy I, bushy II and ameboid cells in PD and control transplanted striatal microglia in absence of LPS challenge. Repeated-measures mixed-effects analysis followed by Sidak’s multiple comparisons test. **C**. Percentage of CD68+ cells in transplanted PD and control striatal microglia in non-inflammatory conditions (no LPS injection). Unpaired t-test. **D**. Integrated density of Iba1 signal of control and PD microglia in non-inflammatory conditions (no LPS injection). Unpaired Mann-Whitney test. **E**. Percentage of p65-positive cells in transplanted PD and control striatal microglia in non-inflammatory conditions (no LPS injection). Unpaired t-test. **F**. Confocal microscopy maximum intensity projection of GFP-labeled control and PD transplanted striatal microglia after LPS injection. Scale bar is 20 µm. **G**. Percentages of ramified, bushy I, bushy II and ameboid cells in PD and control transplanted striatal microglia after LPS injection. Repeated-measures mixed-effects analysis followed by Sidak’s multiple comparisons test. **H**. Confocal maximum intensity projection of Control (left panel) and Parkinson (right panel) GFP-labeled transplanted striatal microglia in a transplanted mouse brain slice after LPS injection, immunostained with CD68 (red). Scale bar is 20 µm. **I**. Percentage of CD68+ cells in transplanted PD and control striatal microglia in pro-inflammatory conditions (after LPS injection). Unpaired t-test. **J**. Confocal plans of control (left panel) and PD (right panel) transplanted striatal microglia immunostained for Iba1. Arrows indicate cell bodies. Scale bar is 5 µm. **K**. Integrated density of Iba1 signal in PD and control transplanted striatal microglia in pro-inflammatory conditions. Unpaired Mann-Whitney test. **L**. Confocal maximum intensity projection of Control (upper panel) and Parkinson (lower panel) GFP-labeled transplanted striatal microglia in a transplanted mouse brain slice after LPS injection, immunostained with p65 (red). Scale bar is 20 µm. Arrows indicate cell bodies. **M**. Percentage of p65+ cells in transplanted control and PD striatal microglia in pro-inflammatory conditions (after LPS injection). Unpaired t-test. N= 3 cell lines per group, 3-5 transplanted mice per cell line. Data is represented as mean±SEM. *: p<0.05; **: p<0.01; ***: p<0.001; ****: p<0.0001.

In non-inflammatory conditions (no LPS injection), striatal PD microglia were largely quiescent, as assessed by low percentages of CD68-positive cells and high percentages of ramified cells (Figures 4B, C and S5A, B). The morphology and percentage of CD68-positive cells were similar in control and PD microglia (Figures 4B, C and S5A, B). No consistent difference in Iba1 staining intensity could be observed between control and PD transplanted microglia (Figures 4D, S5C) and no consistent difference in percentage of p65-positive cells could be found between control and PD microglia (Figures 4E and S5D). Together, these results indicated that, in non-inflammatory conditions, the level of activation of PD and control microglia was similar.

LPS acts as a potent stimulator of microglia, and has been used to study the inflammatory process in the pathogenesis of PD [40]. In previous studies, intraperitoneal injection of LPS was shown to successfully activate human microglia transplanted into the mouse brain [13, 14]. To place transplanted microglia in inflammatory conditions, transplanted mice were injected with LPS at 5 mg/kg. Brains were collected 24 hours after injection. LPS successfully induced activation of control and PD microglia, as assessed by elevated percentages of CD68-positive cells and a shift towards a less ramified morphology compared to non-inflammatory conditions (Figures 4F-I, S6A, B). The distribution of the morphology of transplanted PD microglia was significantly shifted towards less ramified categories compared to control, (Figures 4F, G, S6A), and they displayed slightly but significantly increased percentages of CD68-positive cells (Figures 4H, I, S6B). Additionally, transplanted PD microglia showed increased Iba1 staining intensity (Figures 4J, K, S6C) and higher percentages of p65-positive cells (Figures 4L, M, S6D) after LPS induction. Together, these results indicate that, although in non-inflammatory conditions, transplanted PD and control microglia display similar levels of activation, when placed in pro-inflammatory conditions, transplanted PD microglia show increased pro-inflammatory activation compared to isogenic control.

### Transplanted human A53T-mutant microglia display decreased catalase expression and increased oxidative stress

To confirm decreased expression of catalase in PD transplanted microglia, as assessed by RNA sequencing, on transplanted human microglia at the protein level, we performed immunofluorescence staining against catalase on 20 µm-thick brain slices from transplanted mice at 2 months PI. In line with the RNA sequencing results, we found a strong decrease in catalase signal intensity in PD microglia compared to control in non-inflammatory conditions (Figures 5A, B, S7A). To investigate whether this decrease was associated with increased oxidative stress in PD microglia, we immunostained brain slices from transplanted mice with the DNA oxidative stress marker 8-hydroxy-2’-deoxyguanosine (8-OHdG) at 2-3 months PI. We found strongly increased staining intensity of 8-OHdG, indicating increased oxidative DNA damage in PD transplanted microglia, in both non-inflammatory and pro-inflammatory conditions (Figures 5C-E, S7B,C). Surprisingly, no consistent difference in catalase expression was observed between PD and control in pro-inflammatory conditions (Figure S7D).

**Figure 5.**
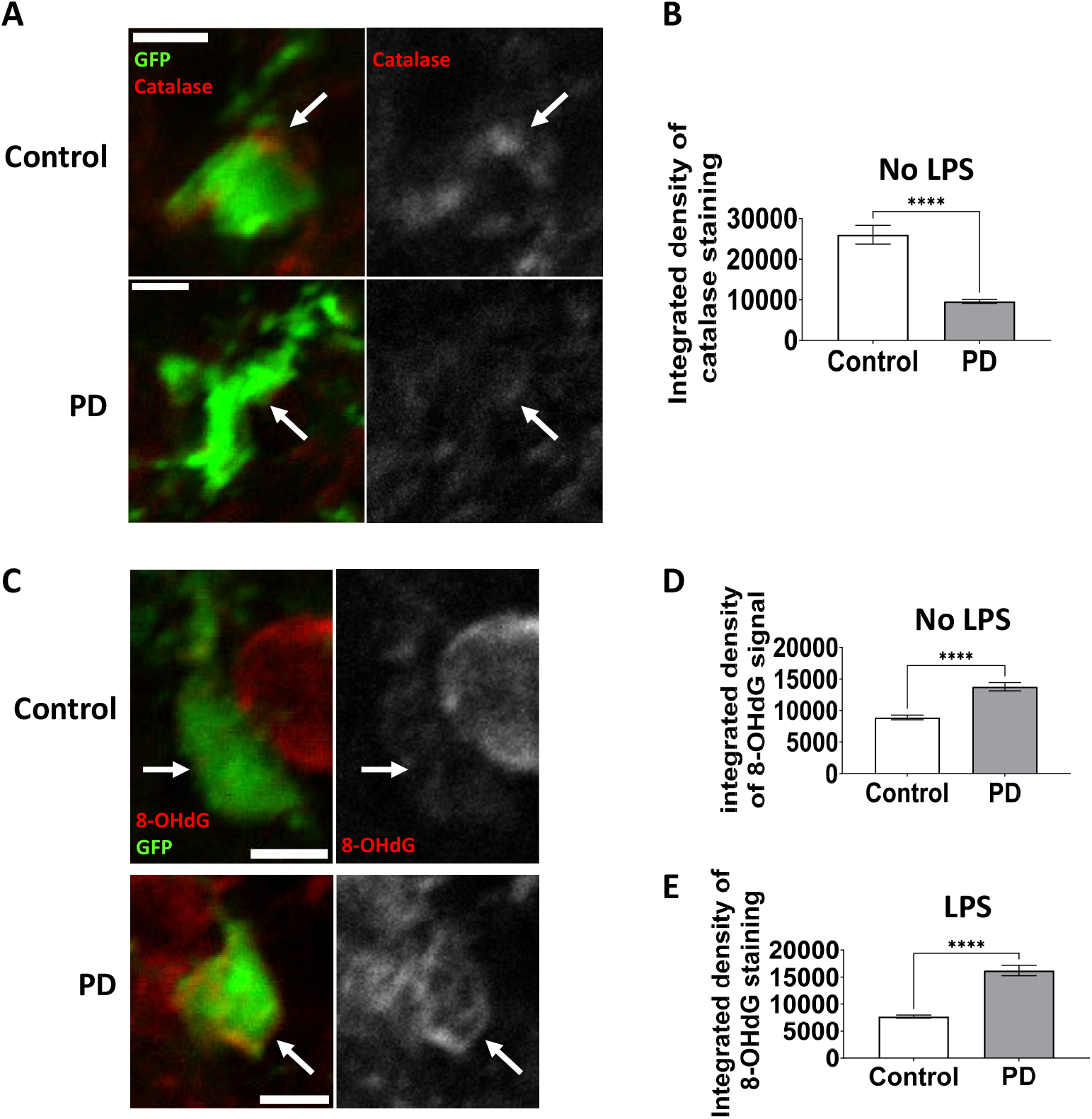
Transplanted PD striatal microglia display strongly decreased catalase expression in non-inflammatory conditions and increased oxidative stress. **A**. Confocal plan of control and PD transplanted striatal microglia at 2 months PI immunostained for catalase (no LPS challenge). Scale bar is 5 µm. Arrows show cell bodies. **B**. Mean integrated density of catalase staining in transplanted control and PD striatal microglia at 2 months PI (no LPS injection). Unpaired Mann-Whitney test. **C**. Confocal plan of control and PD transplanted striatal microglia at 2 months PI immunostained for 8-OHdG (no LPS injection). Scale bar is 5 µm. Arrows show cell bodies. **D**. Mean integrated density of 8-OHdG staining in transplanted control and PD striatal microglia at 2 months PI (no LPS challenge). Unpaired Mann-Whitney test. **E**. Mean integrated density of 8-OHdG staining in transplanted control and PD striatal microglia at 2 months PI after LPS challenge. Unpaired Mann-Whitney test. N= 3 cell lines per group, 3-5 transplanted mice per cell line. Data is represented as mean±SEM.

Because the RNA sequencing data suggested differences in cellular senescence, cell division and apoptosis between PD and control microglia, we immunostained PD and control transplanted microglia for cleaved caspase 3, an apoptosis marker (Figure S8), ki67, a cell division marker (Figure S9) and p16, a marker of cellular senescence (Figure S10). In non-inflammatory conditions and after LPS induction (pro-inflammatory conditions), low percentages of PD and control microglia were stained for these markers. No difference was found between PD and control microglia, indicating that, at least in these conditions, the A53T mutation in α-synuclein did not affect apoptosis, cellular senescence or cell division of microglia.

## Discussion

Here, we assessed the effect of the A53T mutation in α-synuclein in human microglia, using the combination of a traditional *in vitro* culture model, and the transplantation of human MPs into the mouse brain, which allows for differentiation of human microglia in conditions that better mimic the human brain. A53T-mutant microglia displayed gene signatures suggesting an altered activation state, both *in vitro* and when transplanted in a healthy, young animal, compared to isogenic control. In non-inflammatory conditions, transplanted A53T-mutant microglia and isogenic control microglia were largely quiescent and presented similar levels of activation, as assessed by their morphology and activation markers. However, when placed in pro-inflammatory conditions by injecting transplanted mice with LPS, A53T-mutant microglia displayed increased levels of pro-inflammatory activation compared to control. These results show that, even in a young, healthy mouse brain environment, A53T-mutant microglia has an intrinsic higher propensity for a pro-inflammatory activated state upon inflammatory stimulus. In addition, A53T-mutant microglia showed decreased catalase expression and increased oxidative stress. This indicates that A53T-mutant microglia display cell-autonomous phenotypes that may contribute to neuronal damage in early-onset PD (Figure 6).

**Figure 6.**
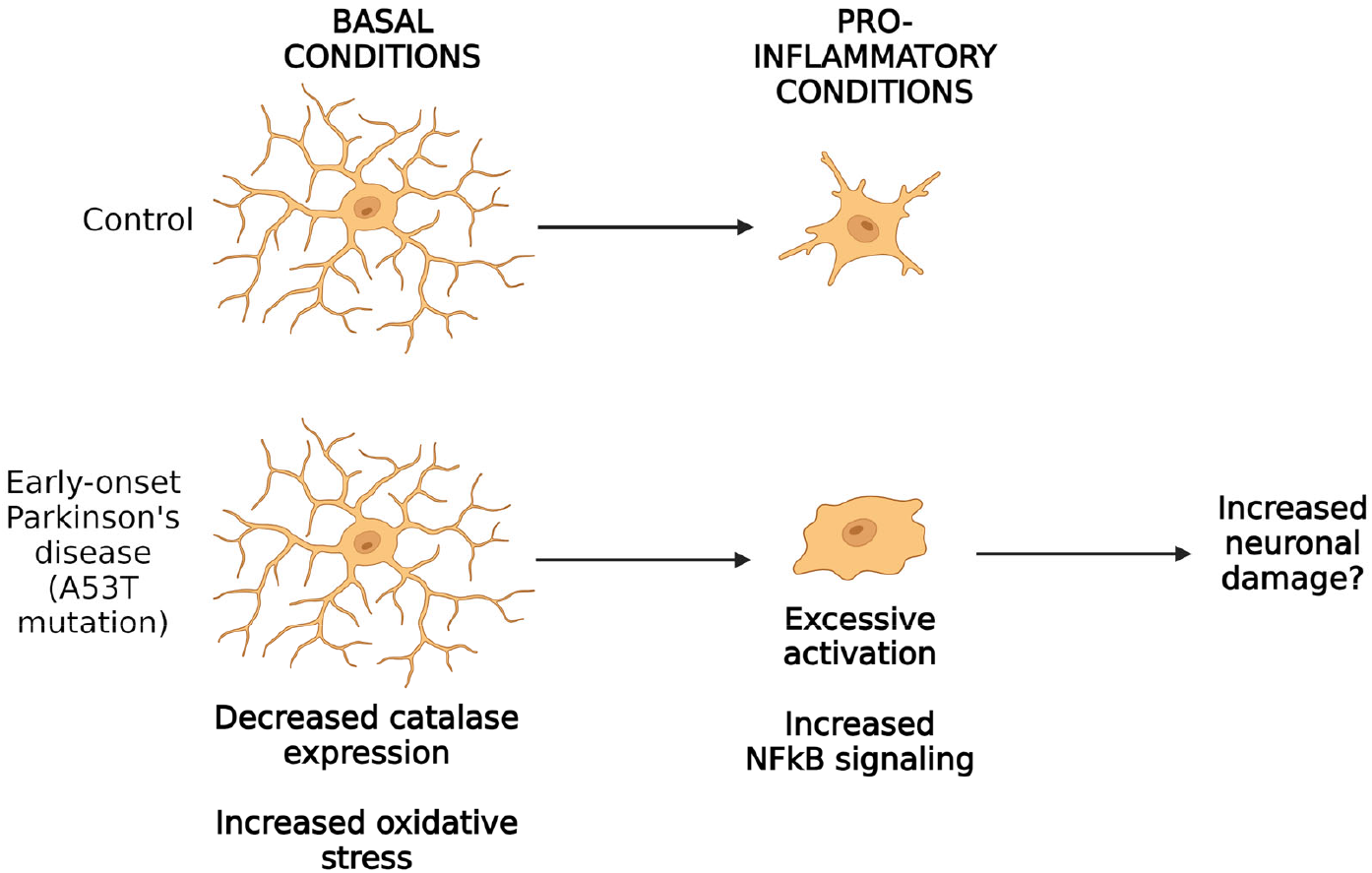
Cell-autonomous disease-relevant phenotypes of A53T-mutant human microglia. In non-inflammatory conditions, A53T-mutant human microglia have decreased expression of catalase and increased oxidative stress. When placed in pro-inflammatory conditions, A53T-mutant human microglia have increased pro-inflammatory activation compared to control, which may trigger increased neuronal damage and worsen neuronal death in early-onset Parkinson’s disease patients.

Mutant α-synuclein inhibits the expression and activity of catalase [41], and catalase activity is decreased in PD patients’ brains and mouse models of PD [41, 42]. Our data showed a strong decrease in catalase expression by A53T-mutant microglia in non-inflammatory conditions. Catalase is involved in the degradation of the ROS H_2_O_2_. Consistent with this, we found increased oxidative stress in PD transplanted microglia, as assessed by oxidative DNA damage, in both pro- and non-inflammatory conditions. Surprisingly however, no consistent differences in catalase expression could be found between PD and control in pro-inflammatory conditions. Although the literature on the link between catalase expression and microglial activation is scarce, one study has pointed to the positive regulation of catalase by the NFκB complex [43]. A potential explanation for the absence of difference in catalase expression between PD and control could be that catalase expression in PD microglia is upregulated in pro-inflammatory conditions, due to the upregulation of the NFκB pathway, as they show increased pro-inflammatory activation in pro-inflammatory conditions.

During activation, microglia generate ROS via NADPH oxidase 2 (NOX2), including H_2_O_2_. These ROS are secreted in the extracellular medium and may damage surrounding cells. They may also enhance microglial pro-inflammatory activation, by activating the NFκB and MAPK signaling pathways [44]. This raises the intriguing possibility that the A53T mutation in α-synuclein leads to increased intracellular concentrations of ROS, increased oxidative stress and subsequent increased microglial activation, triggering a vicious cycle where microglia damages neurons and gets increasingly activated via increased intracellular concentrations of ROS. In a further study, it will be of particular interest to rescue catalase expression in PD microglia and assess whether this leads to decreased oxidative stress, and subsequent decreased microglial activation upon inflammatory stimulus of PD transplanted microglia.

LPS challenge did not trigger increased proliferation of transplanted human microglia. This is surprising, given that previous studies have shown that a single peripheral injection of LPS at 1 mg/kg triggers robust proliferation of mouse microglia, peaking at 24-72h after LPS injection [45, 46]. This lack of proliferation may be due to either decreased sensitivity of transplanted human microglia to LPS compared to mouse microglia, or a longer delay before proliferation of human microglia compared to mouse microglia, as the mice were analyzed 24h after LPS challenge.

The analysis of key microglial genes in transplanted PD and control microglia showed increased expression of several key microglial genes: *CX3CR1, GPR34, P2RY12, P2RY13, SALL1, TGFBR1*. This is consistent with a previous study from our group [13]. The expression of several other genes showed a discrepancy with our previous study. For example, *BIN1* and *SLC2A5* did not show a consistent significant increase in PD and control transplanted microglia. *TMEM119* expression was increased in transplanted PD and control microglia, whereas the previous study showed no increase. This may be due to the fact we used microglia differentiated *in vitro* for one month, whereas the previous study used microglia differentiated *in vitro* for a longer duration, and a different differentiation protocol.

Understanding the impact of A53T mutation on microglia may yield a better understanding of the pathogenesis of PD, as it provides a well-controlled way to assess the dysfunction of these cells in the inherited form of the disease, using isogenic hPSC pairs. Most cases of PD are, however, sporadic, not inherited, and do not involve the A53T mutation. In the future, it will be of interest to extend the significance of our results to the pathogenesis of sporadic PD.

## Acknowledgments and disclosures

This work was supported by grants from the NIH (U19AI131135 and R01MH104610) and by a generous gift from Jim Stone. MK and RJ conceived the idea for this study. MK designed the experiments. MK interpreted the data. BY analyzed the RNA sequencing data. MK, DF, WC, TO, and JS performed the experiments. DSS and KRA contributed to the targeting of the cell lines with GFP at the AAVS1 locus. MK, CGE, WC, and TO analyzed the data. MK and RJ wrote the manuscript with input from all the other authors. We thank Aditya Rathee and Patrick Autissier from the FACS facility of Whitehead Institute for Biomedical Research for FACS and Jennifer Love, Sumeet Gupta and Amanda Chilaka from the Genome Technology Core of Whitehead Institute for RNA sequencing services and consultation. We thank George Bell and Troy W. Whitfield for helpful discussions about statistics. We thank Wendy Salmon, Brandyn Braswell and Cassandra Rogers from the Keck Microscopy Facility of Whitehead Institute for their useful suggestions on confocal microscopy acquisitions. We thank Michael Gallagher, Danielle Tomasello and all the other members of the Jaenisch Lab for discussion and suggestions on the manuscript. Part of this data was presented during the Society for Neuroscience 2022 annual meeting. R.J. is an advisor/co-founder of Fate, Fulcrum, Camp4 and Omega Therapeutics and of Paratus Sciences. All other authors report no biomedical financial interests or potential conflicts of interest.

