## Supplemental figures for "The A53T mutation in α-synuclein enhances pro-inflammatory activation in human microglia"

### Figure S1

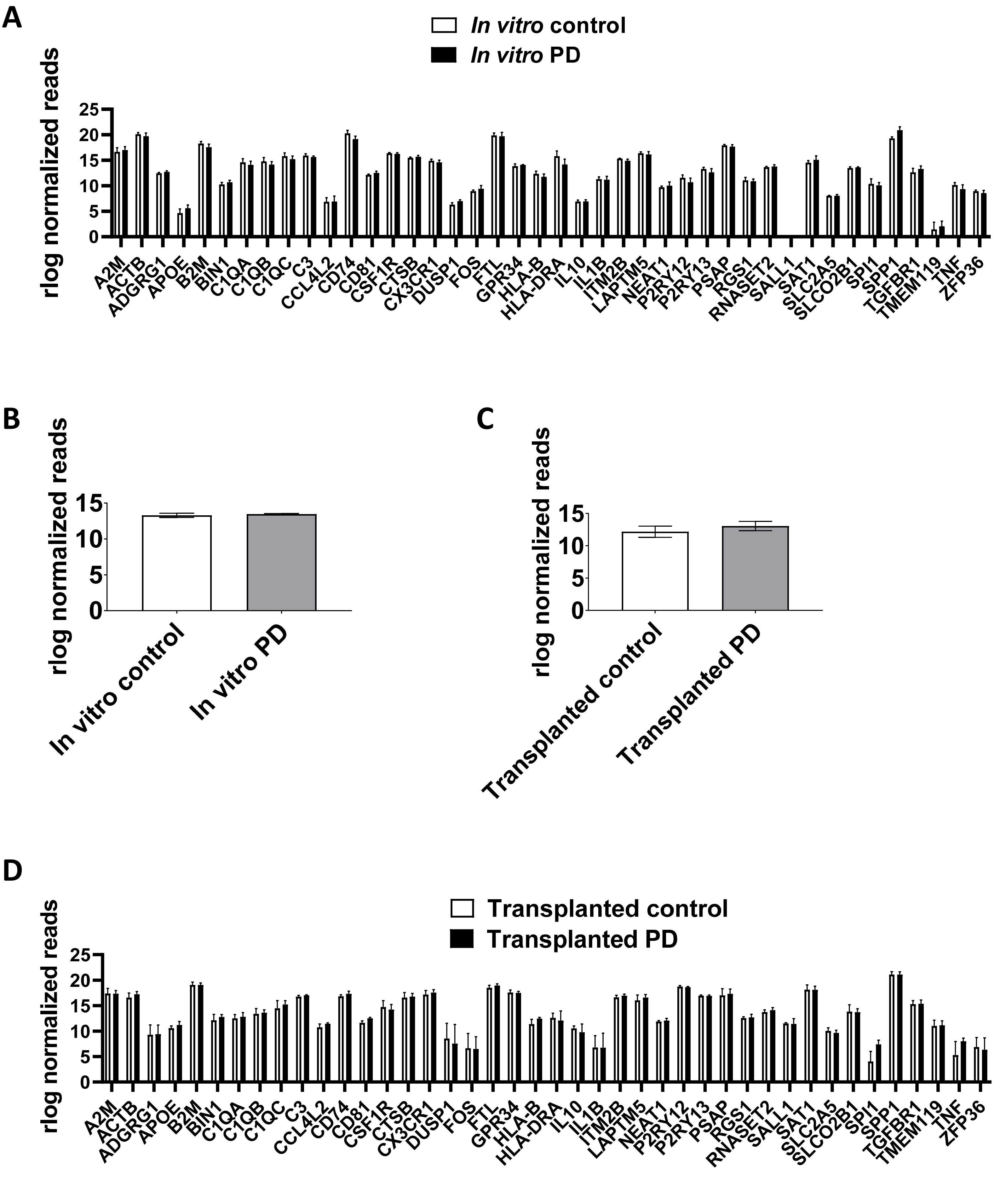
A
In vitro control
In vitro PD
rlog normalized reads
25
20
15
10
5
0
A2M ACTB ADGRG1 APOE B2M BIN1 C1QA C1QB C1QC C3 CCL4L2 CD74 CD81 CSF1R CTSB CX3CR1 DUSP1 FOS FTL GPR34 HLA-B HLA-DRA IL10 IL1B ITM2B LAPTM5 NEAT1 P2RY12 P2RY13 PSAP RGS1 RNASET2 SALL1 SAT1 SLC2A5 SLCO2B1 SPI1 SPP1 TGFBR1 TMEM119 TNF ZFP36
B
rlog normalized reads
15
10
5
0
In vitro control
In vitro PD
C
rlog normalized reads
15
10
5
0
Transplanted control
Transplanted PD
D
Transplanted control
Transplanted PD
rlog normalized reads
25
20
15
10
5
0
A2M ACTB ADGRG1 APOE B2M BIN1 C1QA C1QB C1QC C3 CCL4L2 CD74 CD81 CSF1R CTSB CX3CR1 DUSP1 FOS FTL GPR34 HLA-B HLA-DRA IL10 IL1B ITM2B LAPTM5 NEAT1 P2RY12 P2RY13 PSAP RGS1 RNASET2 SALL1 SAT1 SLC2A5 SLCO2B1 SPI1 SPP1 TGFBR1 TMEM119 TNF ZFP36

### Figure S2

**A**

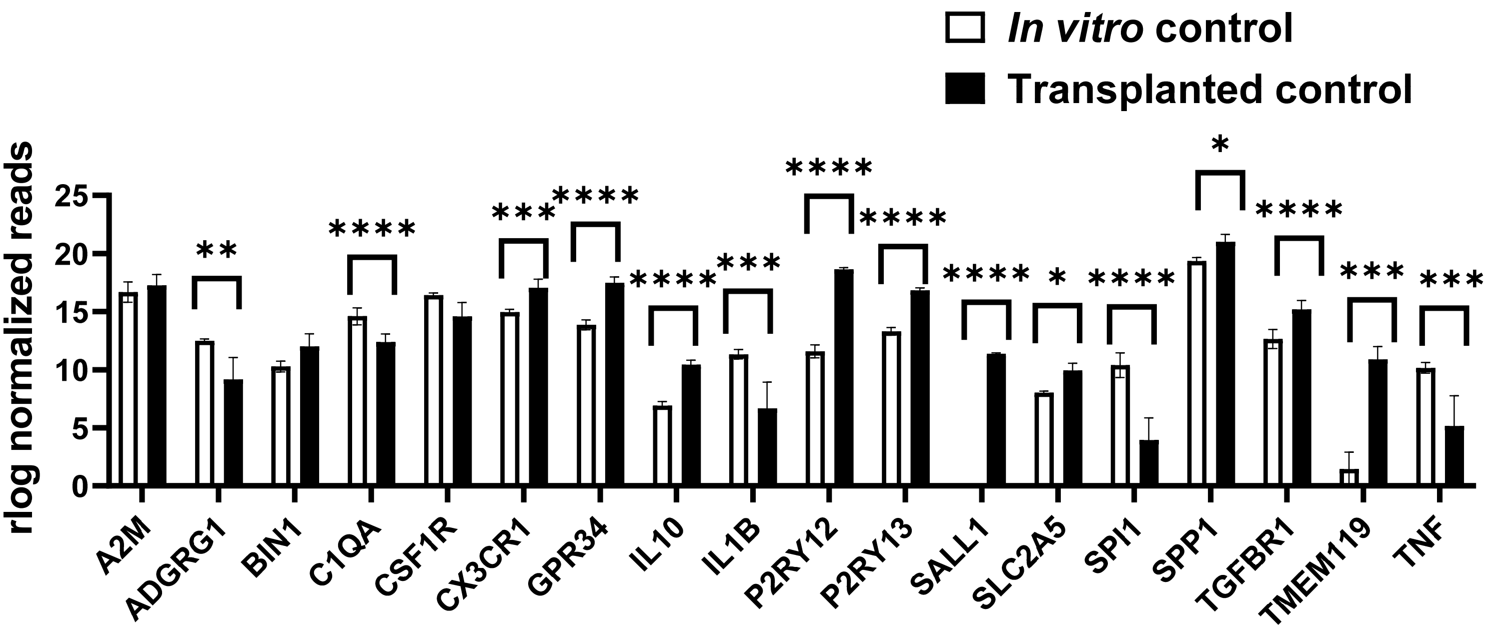

In vitro control
Transplanted control
rlog normalized reads
25
20
15
10
5
0
**
****
***
****
****
***
****
****
****
*
****
*
****
***
***
A2M
ADGRG1
BIN1
C1QA
CSF1R
CX3CR1
GPR34
IL10
IL1B
P2RY12
P2RY13
SALL1
SLC2A5
SPI1
SPP1
TGFBR1
TMEM119
TNF


**B**

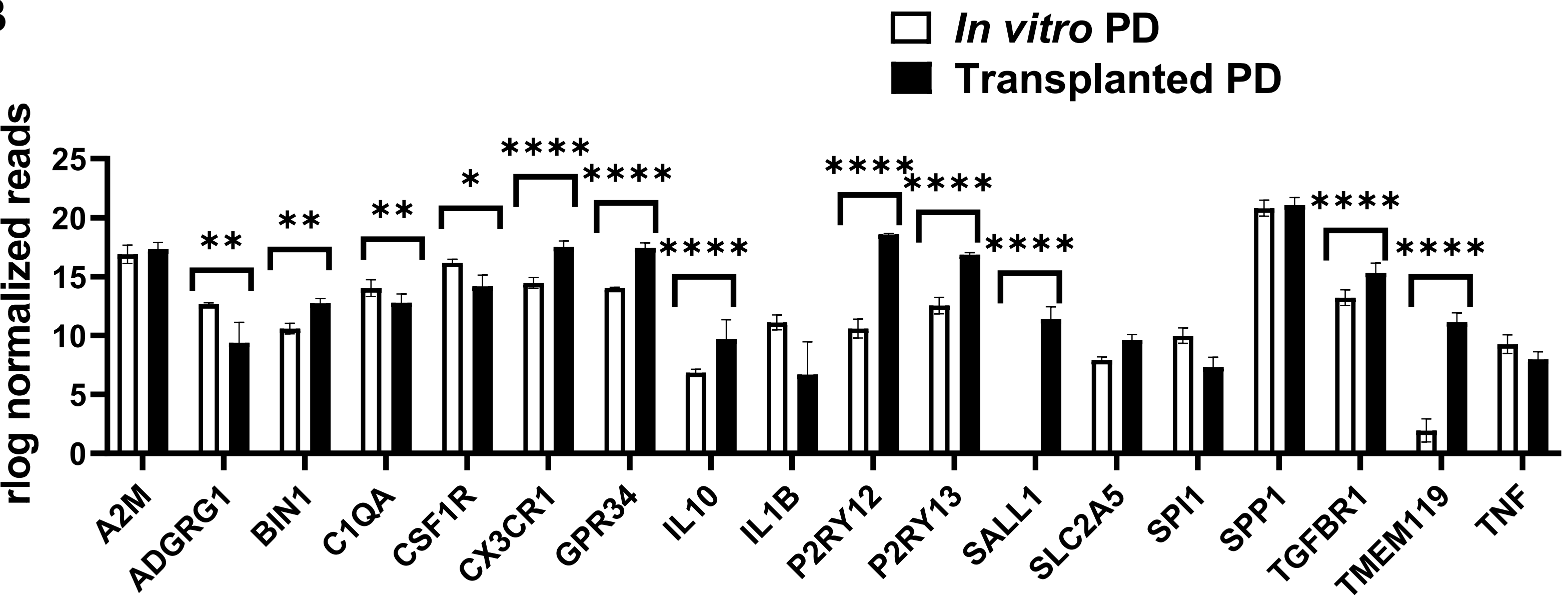

In vitro PD
Transplanted PD
rlog normalized reads
25
20
15
10
5
0
**
**
**
*
****
****
****
****
****
****
****
****
A2M
ADGRG1
BIN1
C1QA
CSF1R
CX3CR1
GPR34
IL10
IL1B
P2RY12
P2RY13
SALL1
SLC2A5
SPI1
SPP1
TGFBR1
TMEM119
TNF

### Figure S3

**A**

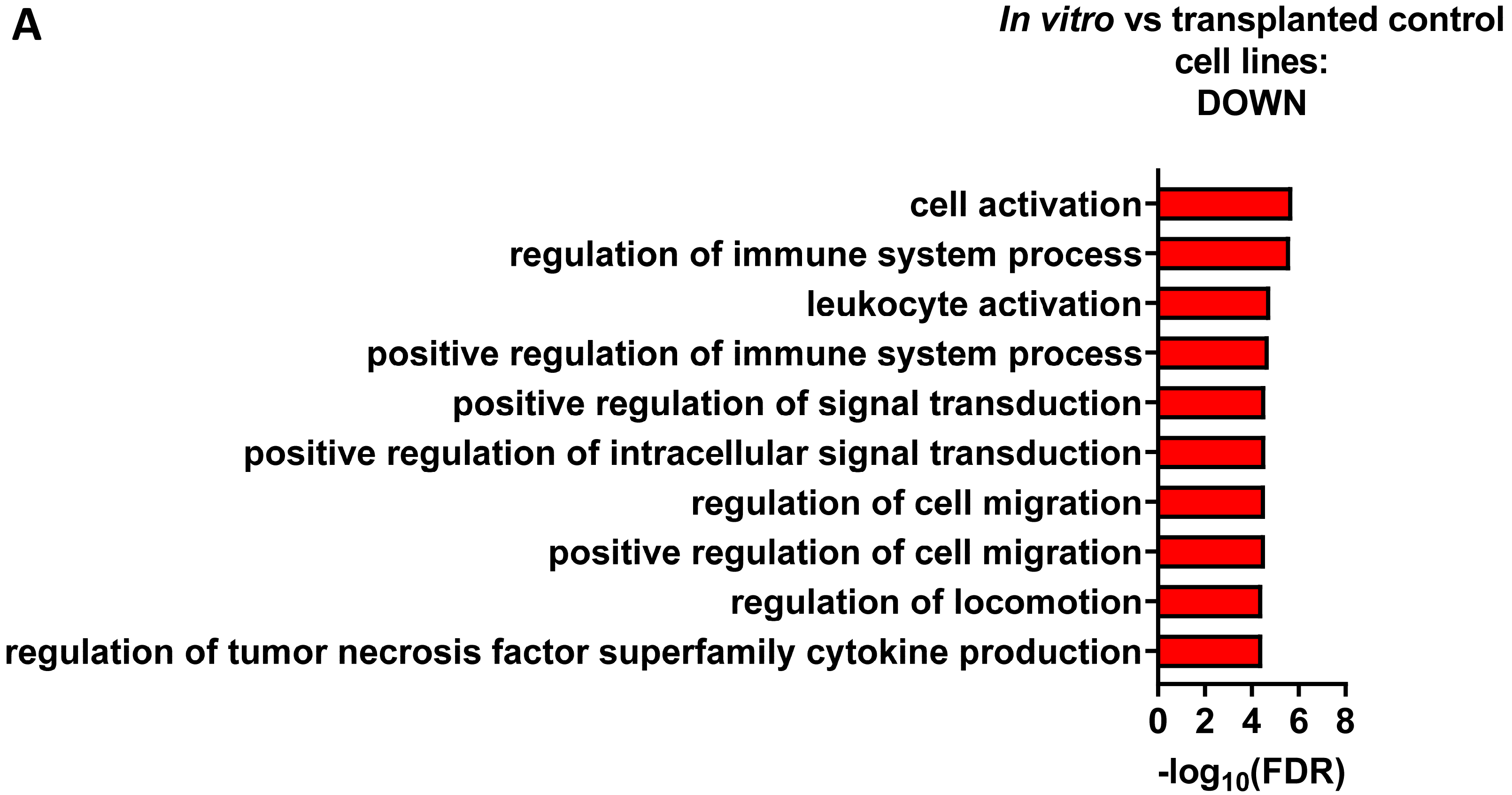
In vitro vs transplanted control cell lines:
DOWN
cell activation
regulation of immune system process
leukocyte activation
positive regulation of immune system process
positive regulation of signal transduction
positive regulation of intracellular signal transduction
regulation of cell migration
positive regulation of cell migration
regulation of locomotion
regulation of tumor necrosis factor superfamily cytokine production
0
2
4
6
8
-log10(FDR)

**B**

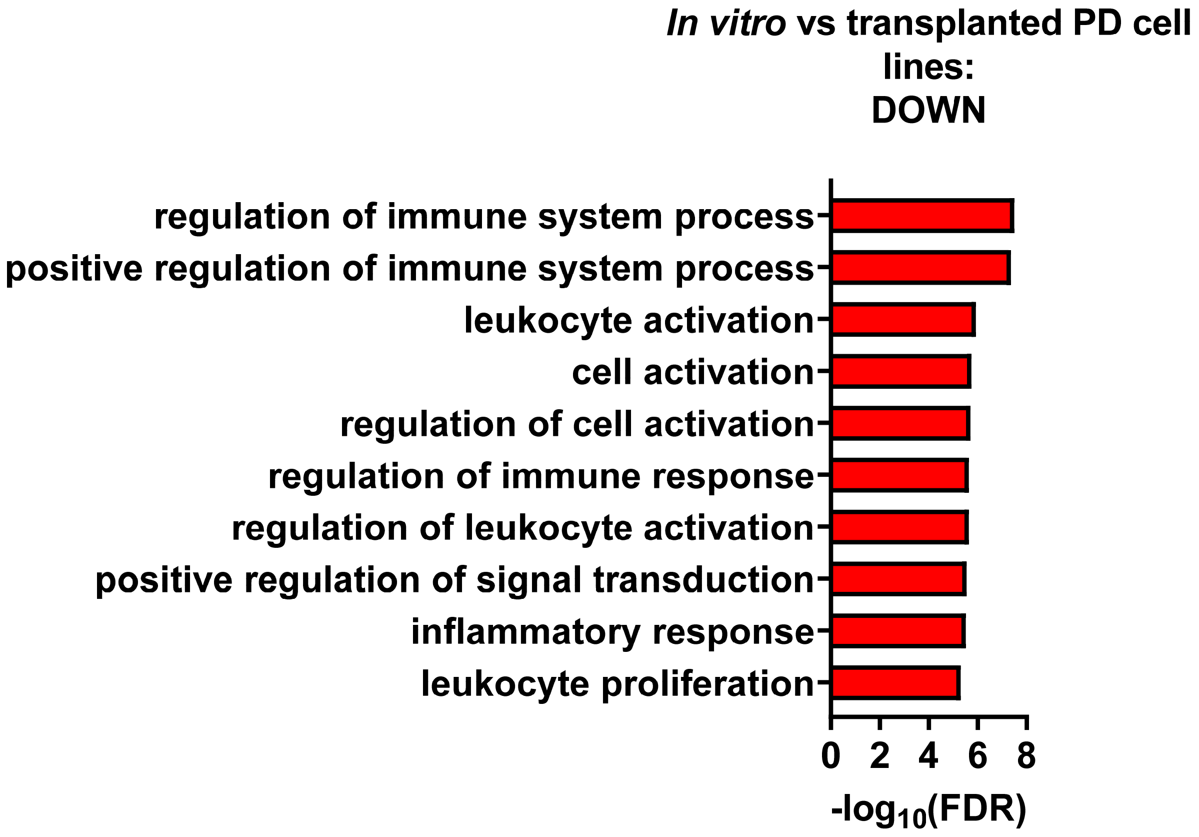
In vitro vs transplanted PD cell lines:
DOWN
regulation of immune system process
positive regulation of immune system process
leukocyte activation
cell activation
regulation of cell activation
regulation of immune response
regulation of leukocyte activation
positive regulation of signal transduction
inflammatory response
leukocyte proliferation
0
2
4
6
8
-log10(FDR)

### Figure S4

A

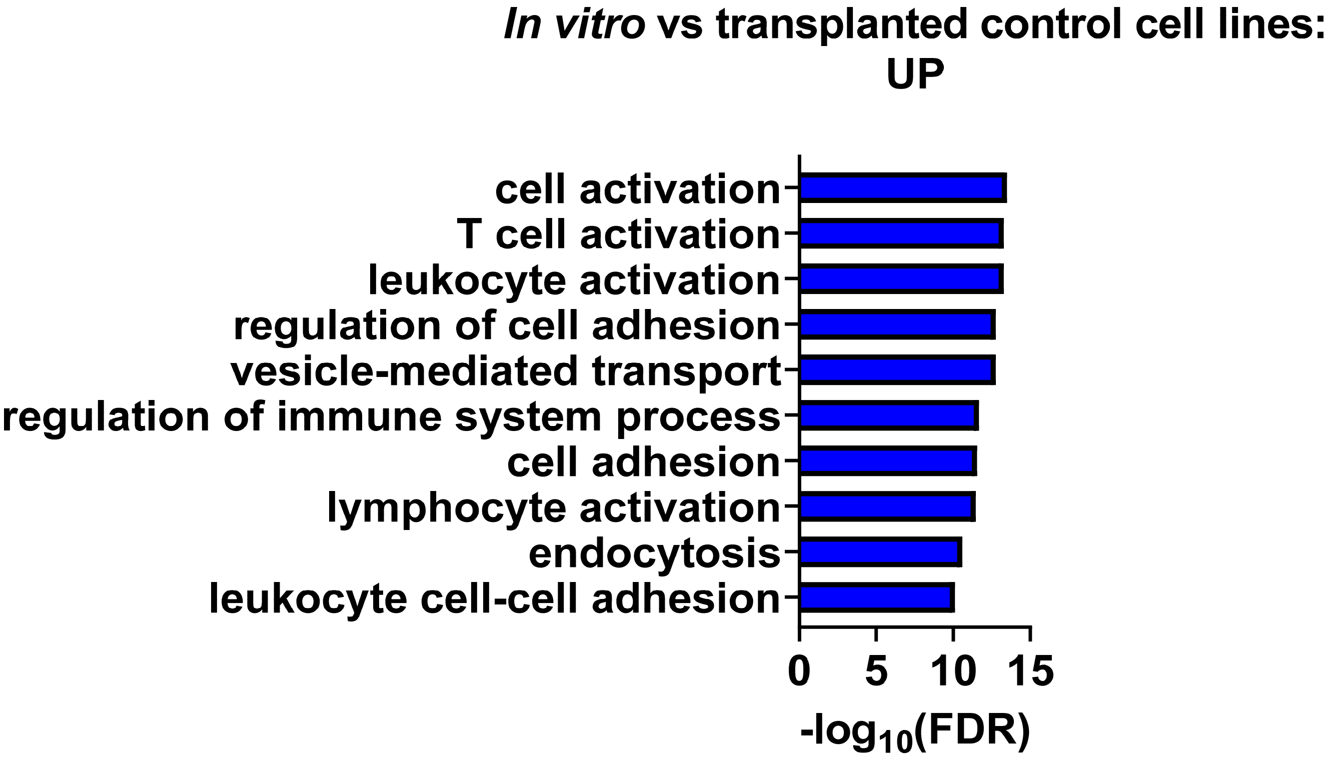
*In vitro* vs transplanted control cell lines:
UP
cell activation
T cell activation
leukocyte activation
regulation of cell adhesion
vesicle-mediated transport
regulation of immune system process
cell adhesion
lymphocyte activation
endocytosis
leukocyte cell-cell adhesion
0
5
10
15
-log10(FDR)

B

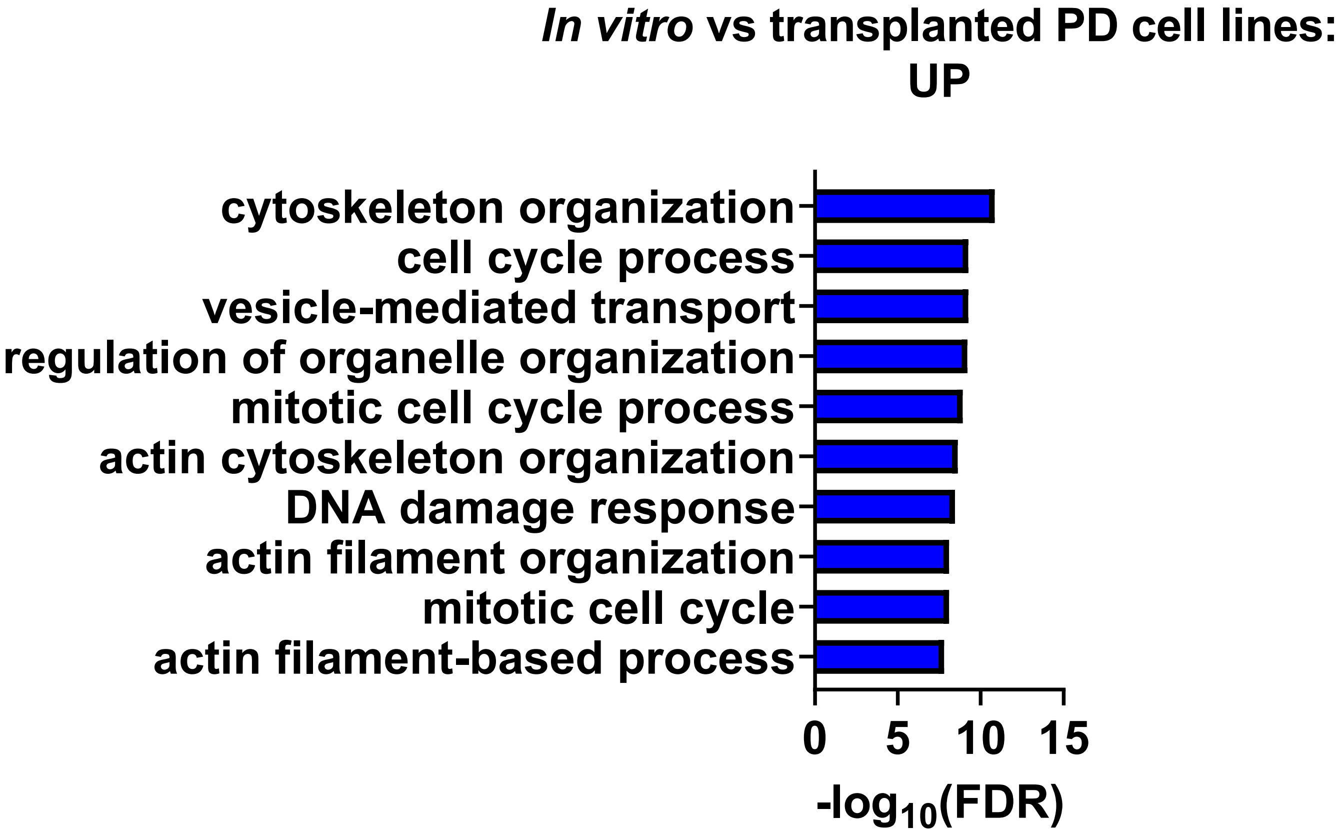
*In vitro* vs transplanted PD cell lines:
UP
cytoskeleton organization
cell cycle process
vesicle-mediated transport
regulation of organelle organization
mitotic cell cycle process
actin cytoskeleton organization
DNA damage response
actin filament organization
mitotic cell cycle
actin filament-based process
0
5
10
15
-log10(FDR)

### Figure S5

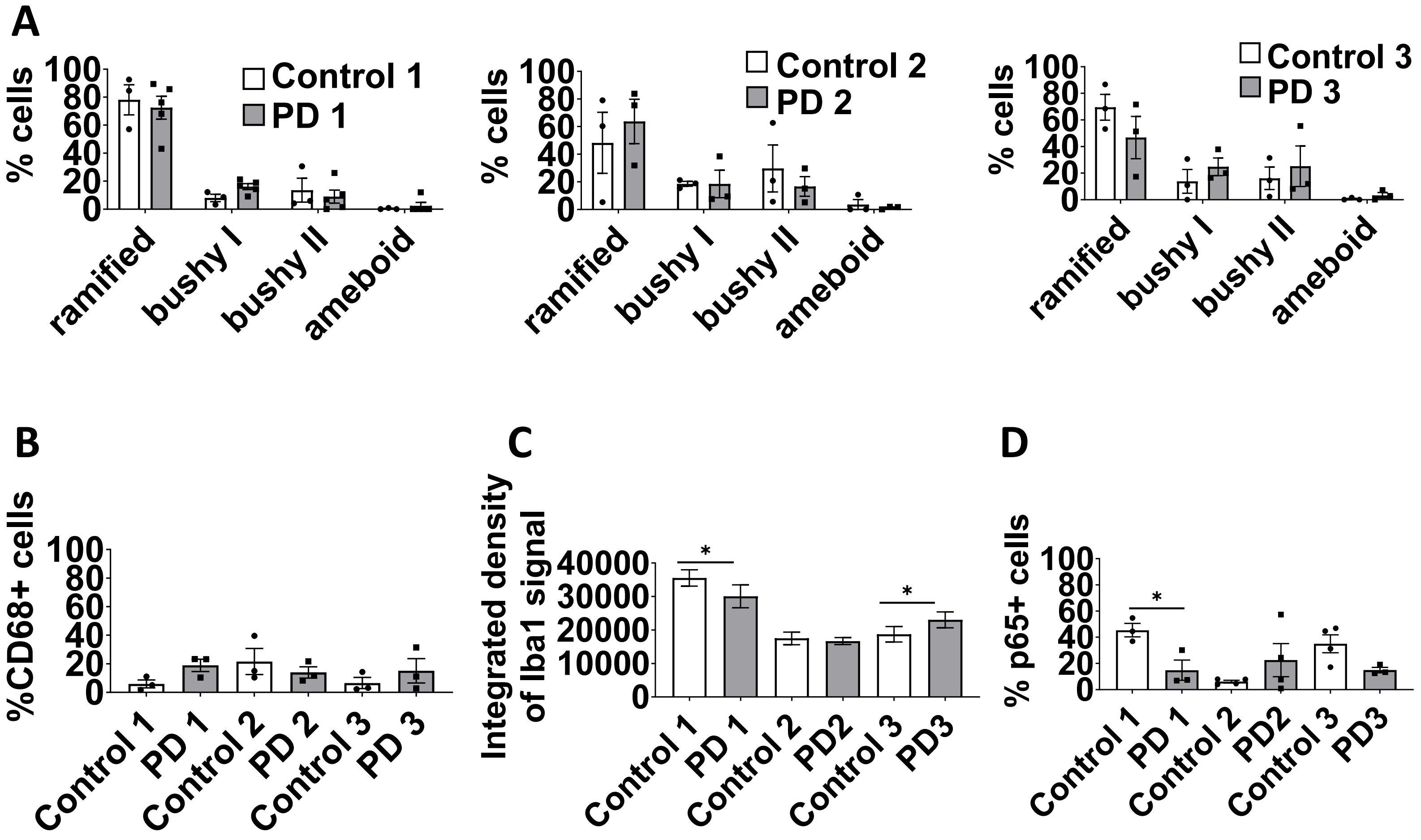
A
% cells
Control 1
PD 1
ramified
bushy I
bushy II
ameboid
Control 2
PD 2
Control 3
PD 3
B
%CD68+ cells
Control 1
PD 1
Control 2
PD 2
Control 3
PD 3
C
Integrated density of Iba1 signal
PD2
PD3
D
% p65+ cells

### Figure S6

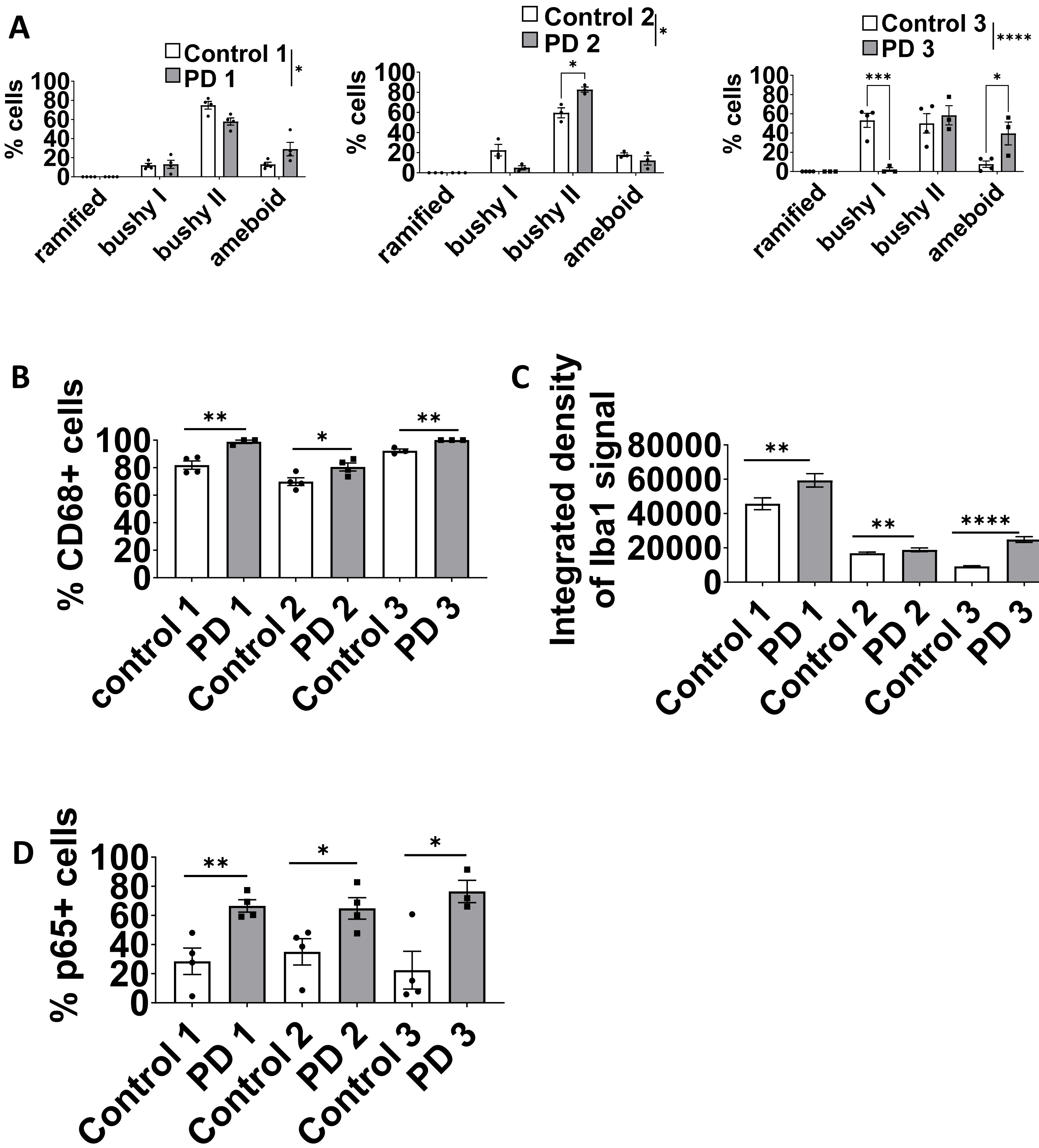
A
Control 1
PD 1
*
% cells
100
80
60
40
20
0
ramified
bushy I
bushy II
ameboid
Control 2
PD 2
*
Control 3
PD 3
****
***
B
% CD68+ cells
**
control 1
PD 1
Control 2
PD 2
Control 3
PD 3
C
Integrated density of Iba1 signal
80000
60000
40000
20000
Control 1
D
% p65+ cells

### Figure S7

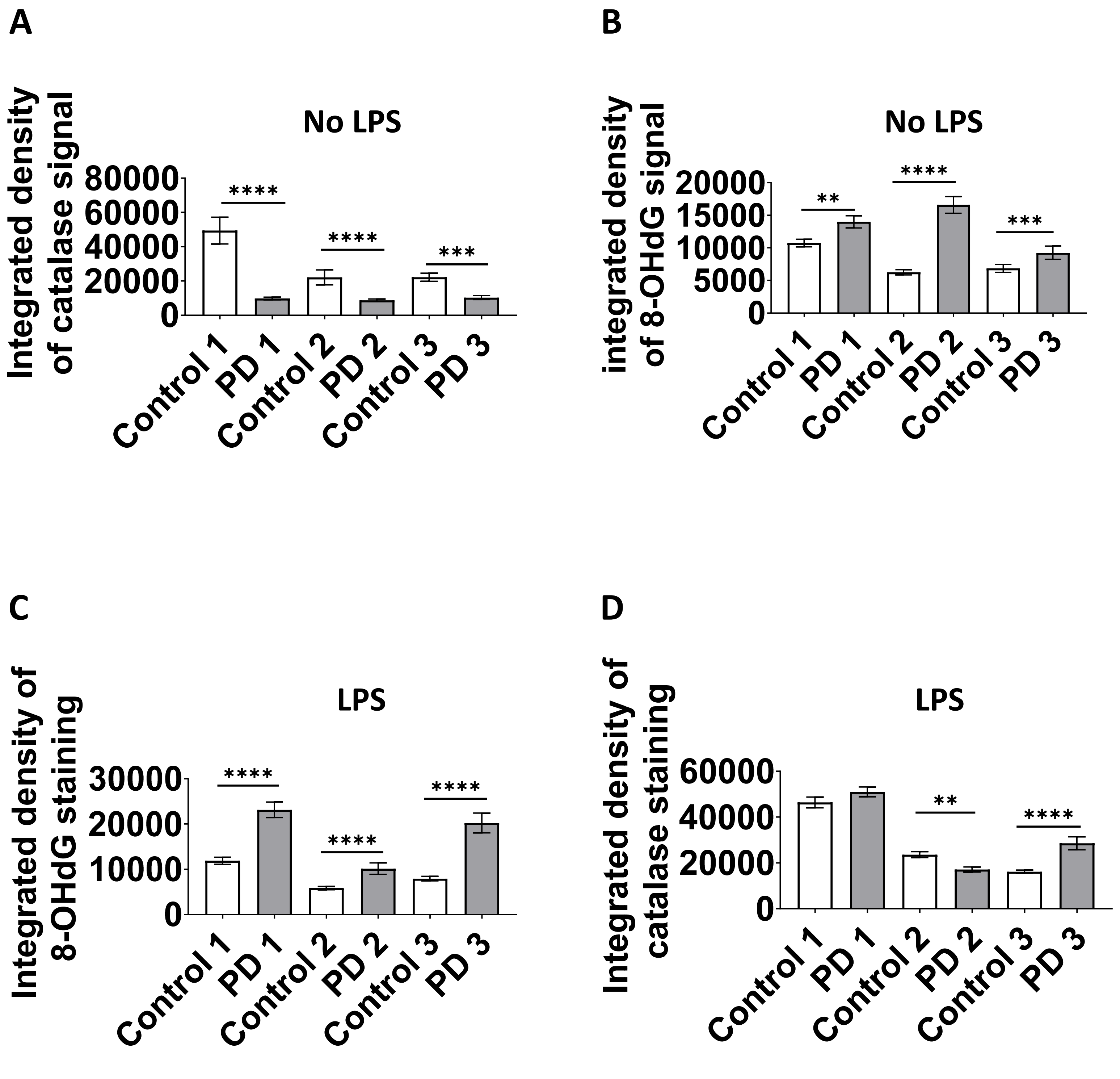
A
No LPS
Integrated density of catalase signal
80000
60000
40000
20000
0
****
****
***
Control 1
PD 1
Control 2
PD 2
Control 3
PD 3
B
No LPS
integrated density of 8-OHdG signal
20000
15000
10000
5000
0
**
****
***
Control 1
PD 1
Control 2
PD 2
Control 3
PD 3
C
LPS
Integrated density of 8-OHdG staining
30000
20000
10000
0
****
****
****
Control 1
PD 1
Control 2
PD 2
Control 3
PD 3
D
LPS
Integrated density of catalase staining
60000
40000
20000
0
**
****
Control 1
PD 1
Control 2
PD 2
Control 3
PD 3

### Figure S8

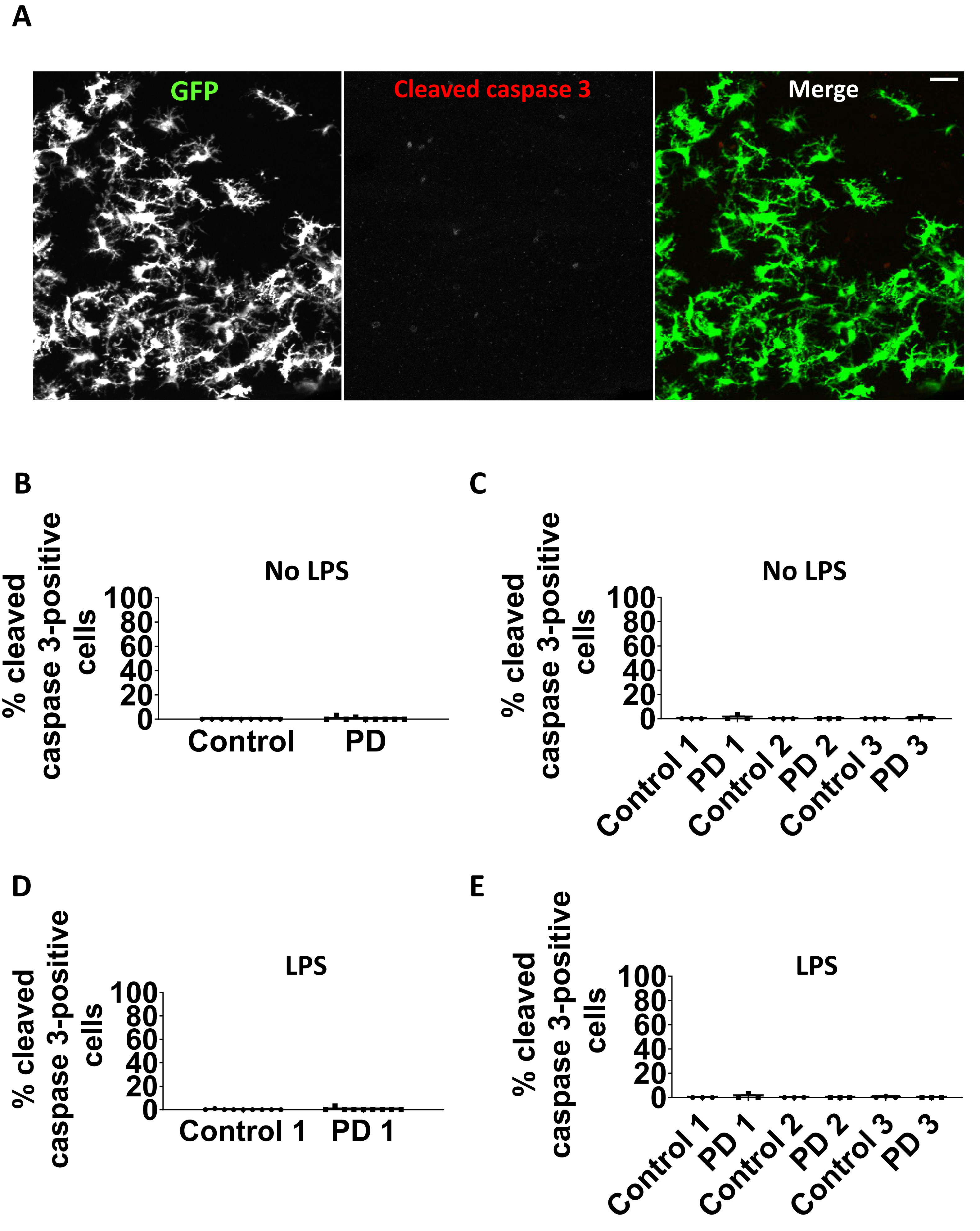
A
GFP
Cleaved caspase 3
Merge
B
No LPS
% cleaved caspase 3-positive cells
100
80
60
40
20
0
Control
PD
C
No LPS
% cleaved caspase 3-positive cells
Control 1
PD 1
Control 2
PD 2
Control 3
PD 3
D
LPS
% cleaved caspase 3-positive cells
Control 1
PD 1
E
LPS
% cleaved caspase 3-positive cells
Control 1
PD 1
Control 2
PD 2
Control 3
PD 3

### Figure S9

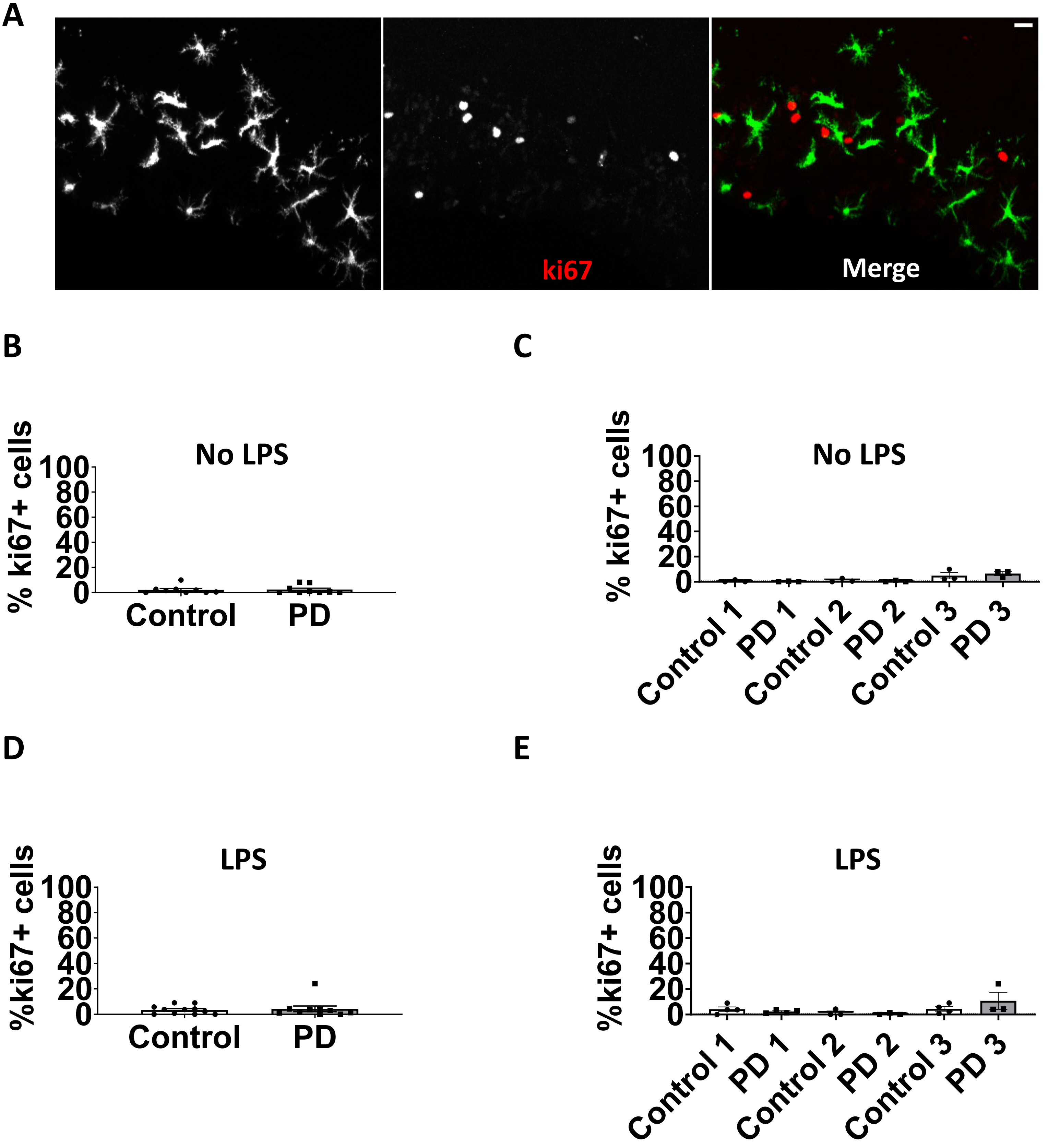
A
ki67
Merge
B
No LPS
% ki67+ cells
100
80
60
40
20
0
Control
PD
C
No LPS
% ki67+ cells
100
80
60
40
20
0
Control 1
PD 1
Control 2
PD 2
Control 3
PD 3
D
LPS
%ki67+ cells
100
80
60
40
20
0
Control
PD
E
LPS
%ki67+ cells
100
80
60
40
20
0
Control 1
PD 1
Control 2
PD 2
Control 3
PD 3

### Figure S10

A

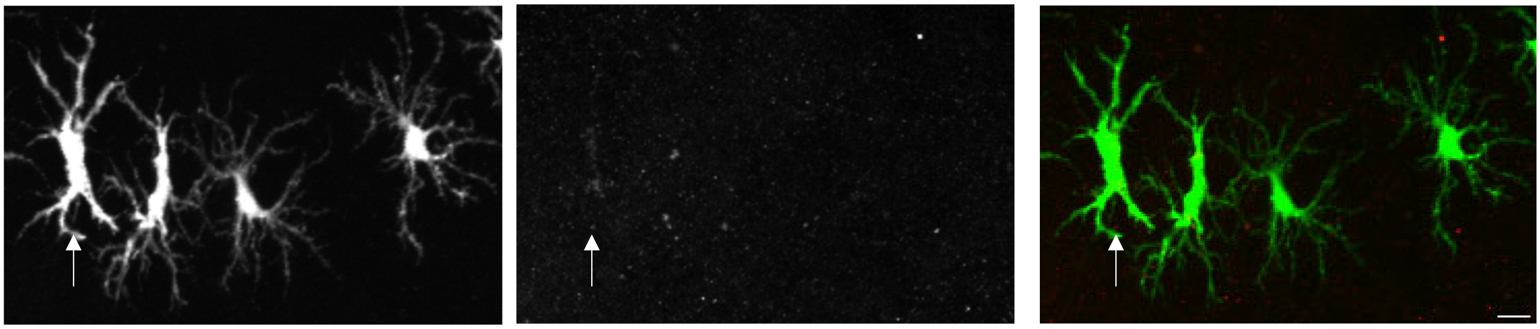

B

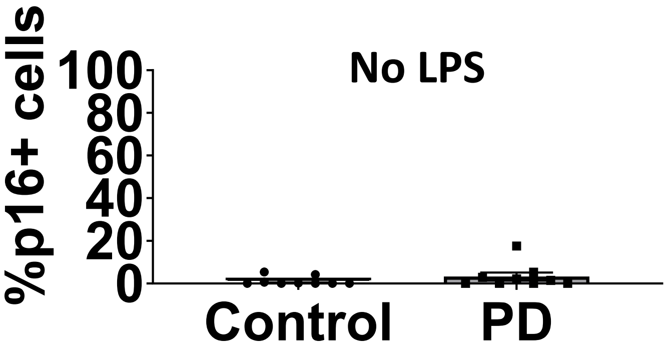


C

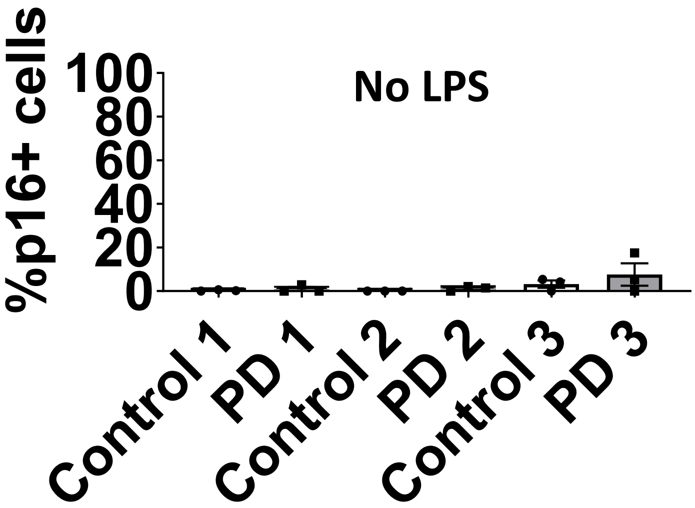


D

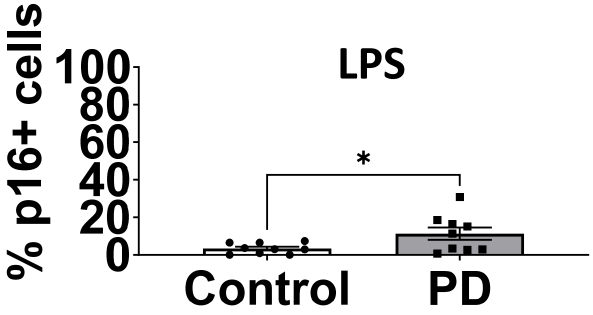


E

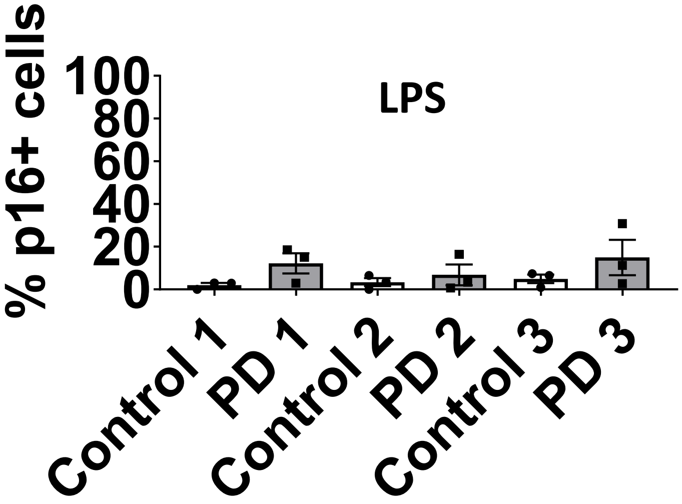
