## Supplementary material for "The A53T mutation in α-synuclein enhances pro-inflammatory activation in human microglia": Tables

### Table 1

| Pair | Isogenic control | Parkinson |
|---|---|---|
| **Control 1/ PD1** | WIBR-hiPS-SNCA $^{A53T-Corr}$: Induced pluripotent stem cell line with corrected A53T mutation in SNCA gene | WIBR-hiPS-SNCA$^{A53T}$ : Early-onset Parkinson patient-derived induced pluripotent stem cell line with heterozygous A53T mutation in SNCA gene |
| **Control 2/ PD 2** | WIBR3: Human embryonic stem cell line | WIBR3-SNCA$^{A53T/WT}$: Embryonic stem cell line WIBR3 with inserted heterozygous A53T mutation in SNCA gene |
| **Control 3/ PD 3** | BGO1: Human embryonic stem cell line | BGO1-SNCA$^{A53T/WT}$: Embryonic stem cell line BGO1 with inserted heterozygous A53T mutation in SNCA gene |

### Table 2

| Antigen | Species | Source | Identifier |
|---|---|---|---|
| 8-Hydroxy-2'-deoxyguanosine (8 OhdG) | Mouse | R and D systems | 4354-MC-050 |
| Catalase | Mouse | Thermofisher | MA42573 |
| CD68 | Mouse | Life Technologies | MA513324 |
| Cleaved caspase 3 | Rabbit | Cell Signaling | 9661T |
| GFP | Chicken | AVES Labs | GFP-1020 |
| Iba1 | Goat | Abcam | ab5076 |
| ki67 | Rabbit | Thermofisher | MA514520 |
| P2RY12 | Rabbit | Sigma-Aldrich | HPA014518 |
| p16 | Mouse | Life technologies | CF500036 |

### Table 3

| Secondary antibody | Source | Identifier |
|---|---|---|
| Donkey Alexa fluor 488 anti-chicken | Jackson ImmunoResearch | 703-545-155 |
| Donkey Alexa fluor 555 anti-goat | Life Technologies | A21432 |
| Donkey Alexa fluor 594 anti-mouse | Life Technologies | A21203 |
| Donkey Alexa fluor 594 anti-rabbit | Life Technologies | A21207 |
| Donkey Alexa fluor 647 anti-mouse | Life Technologies | A31571 |
| Donkey Alexa fluor 647 anti-rabbit | Life Technologies | A31573 |

### Table 4

| Cell line | % Iba1+ P2RY12+ cells |
|---|---|
| **Control 1** | 86.±3.2 |
| **PD 1** | 94±1.6 |
| **Control 2** | 98±0.0 |
| **PD 2** | 98±1.1 |
| **Control 3** | 81±0.7 |
| **PD3** | 95±1.7 |

### Table 5

| Gene symbol | Gene name | Gene ontology | $Log_2(FC)$ |
|---|---|---|---|
| **LBH** | LBH regulator of WNT signaling pathway | Negative regulation of transcription | -1.6 |
| **FCGR2B** | Fc gamma receptor IIb | **Inhibitory receptor of immune cells; inhibits microglia activation** | -1.5 |
| **NOTCH4** | notch receptor 4 | **Anti-inflammatory activity in activated macrophages** | -1.4 |
| **CD200R1** | CD200 receptor 1 | **Inhibition of the secretion of pro-inflammatory molecules by microglia; stimulation results in neuroprotection in a model of PD** | -1.4 |
| **BASP1** | brain abundant membrane attached signal protein 1 | Membrane-bound protein | -1.3 |
| **MCOLN2** | mucolipin TRP cation channel 2 | **Possible role in innate immune response** | -1.3 |
| **CDKN1C** | cyclin dependent kinase inhibitor 1C | Negative regulator of cell proliferation | -1.3 |
| **CA2** | carbonic anhydrase 2 | **Increased in aging and neurodegeneration** | -1.3 |
| **SEMA3C** | semaphorin 3C | **Induces apoptosis of activated pro-inflammatory microglia** | -1.3 |
| **CKB** | creatine kinase B | **Suppresses ferroptosis; ferroptosis may contribute to neurodegeneration** | -1.2 |

### Table 6

| Gene symbol | Gene name | Gene ontology | $Log_2(FC)$ |
|---|---|---|---|
| **RETN** | resistin | **Involved in immune defense** | 2.0 |
| **MCEMP1** | mast cell expressed membrane protein 1 | **Predicted to be involved in regulating immune response** | 1.7 |
| **PLTP** | phospholipid transfer protein | **Increased in Alzheimer's disease; Deletion increases microglial phagocytosis and reduces cerebral amyloid ß deposition in a mouse model of Alzheimer's disease** | 1.4 |
| **SLC11A1** | solute carrier family 11 member 1 | **Involved in protection against ROS in macrophages; associated with inflammatory diseases** | 1.4 |
| **SLC35F2** | solute carrier family 35 member F2 | Predicted to enable transmembrane transporter activity | 1.4 |
| **RIN2** | Ras and Rab interactor 2 | Involved in membrane trafficking in the early endocytic pathway | 1.3 |
| **MAPK13** | mitogen-activated protein kinase 13 | **Contributes to inflammation by promoting cytokine release by microglia** | 1.3 |
| **UCN** | urocortin | **Inhibits microglia activation** | 1.3 |
| **MBOAT1** | membrane bound O-acyltransferase domain containing 1 | Transfers organic compounds to hydroxyl groups of protein targets in membranes | 1.3 |
| **MSR1** | macrophage scavenger receptor 1 | **Secretion of pro-inflammatory cytokines by macrophages; Involved in the uptake and clearance of soluble amyloid ß in Alzheimer's disease by microglia** | 1.3 |

### Table 7

| Cell line | % Iba1+ P2RY12+ cells |
|---|---|
| **Control 1** | 93±4.5 |
| **PD 1** | 100±0.0 |
| **Control 2** | 94±3.5 |
| **PD 2** | 97±2.9 |
| **Control 3** | 100±0.0 |
| **PD3** | 89±0.7 |

### Table 8

| Gene symbol | Gene name | Gene ontology | $Log_2$(FC) |
|---|---|---|---|
| **CAT** | catalase | **Activity decreased in Parkinson patient brains**<br>**Mitigates oxidative stress by breaking down ROS** | -3.8 |
| **ZNF721** | zinc finger protein 721 | Transcription factor | -1.7 |
| **ANLN** | anillin, actin binding protein | Cell growth and migration, cytokinesis | -1.5 |
| **UHRF1BP1** | UHRF1 binding protein 1 | Enables histone deacetylase binding activity and identical protein binding activity | -1.5 |
| **GPR137B** | G protein-coupled receptor 137B | positive regulation of TORC1 signaling; positive regulation of protein localization to lysosome and lysosome morphology and regulation of GTPase activity | -1.5 |
| **CDCP1** | CUB domain containing protein 1 | Involved in cell adhesion, cell matrix association, and T cell activation, migration and chemotaxis | -1.4 |
| **NUP58** | nucleoporin 58 | Component of the nuclear pore complex | -1.4 |
| **KIF27** | kinesin family member 27 | Role in Hedgehog signaling pathway? | -1.4 |
| **SLFN5** | schlafen family member 5 | Predicted to be involved in cell differentiation | -1.3 |
| **PLAU** | plasminogen activator, urokinase | May be associated with late-onset Alzheimer's disease | -1.3 |

### Table 9

| Gene symbol | Gene name | Gene ontology | $Log_2(FC)$ |
|---|---|---|---|
| **INTU** | inturned planar cell polarity protein | key role in ciliogenesis and embryonic development | 2.1 |
| **ZNF613** | zinc finger protein 613 | Transcription factor | 1.5 |
| **OPHN1** | oligophrenin 1 | Implicated in synaptic function | 1.5 |
| **RAB4A** | RAB4A, member RAS oncogene family | associated with early endosomes and is involved in their sorting and recycling; involved in Alzheimer's disease | 1.3 |
| **B3GNT2** | UDP-GlcNAc:betaGal beta-1,3-N-acetylglucosaminyltransferase 2 | Transmembrane protein | 1.2 |
| **RBMS3** | RNA binding motif single stranded interacting protein 3 | **implicated in DNA replication, gene transcription, cell cycle progression and apoptosis; linked to Alzheimer's disease and to motor complications in PD** | 1.2 |
| **CCDC125** | coiled-coil domain containing 125 | Negative regulation of cell motility | 1.2 |
| **KLHL24** | kelch like family member 24 | ubiquitin ligase substrate receptor | 1.1 |
| **TBX15** | T-box transcription factor 15 | Transcription factor regulating developmental processes | 1.1 |
