## Supplemental figure and table legends for "The A53T mutation in α-synuclein enhances pro-inflammatory activation in human microglia"

### Supplemental figures

**Figure S1. Control and PD microglia cultured *in vitro* or transplanted into the mouse brain display similar levels of expression of key microglial genes, indicating similar levels of differentiation. A.** Expression of key microglial genes in control and PD human microglia differentiated *in vitro* for 29 days. **B.** Expression of SNCA in control and PD human microglia differentiated *in vitro* for 29 days. **C.** Expression of SNCA in transplanted control and PD human microglia at 2 months post-transplantation. **D.** Expression of key microglial genes in transplanted control and PD human microglia at 2 months post-transplantation. Values are from RNA sequencing experiments with read counts normalized by library size. Significant differences in the level of expression of each gene were defined as a log2-fold difference strictly superior to 1 or strictly inferior to 1 and $p_{adjusted}<0.05$. n=3 cell lines per group, 3 biological experiments for each cell line.

**Figure S2. Transplanted human microglia displays increased expression of a number of key microglial genes compared to *in vitro* differentiated microglia. A.** Expression of key microglial genes in control human microglia differentiated *in vitro* for 29 days (*In vitro* control) or at 2 months post-transplantation into the brain of mice (Transplanted control). **B.** Expression of key microglial genes in PD human microglia differentiated *in vitro* for 29 days (*In vitro* control) or at 2 months post-transplantation into the brain of mice (Transplanted control). Values are from RNA sequencing experiments with read counts normalized by library size. Significant differences in the level of expression of each gene were defined as a log2-fold difference strictly superior to 1 or strictly inferior to 1 and $p_{adjusted}<0.05$. n=3 cell lines per group, 3 biological experiments for each cell line or 2-3 transplanted brains per cell line.

**Figure S3. Downregulated gene pathways in human microglia differentiated *in vitro* vs transplanted human microglia. A.** Ten most downregulated gene pathways in control *in vitro* differentiated microglia compared to control transplanted microglia, as assessed by gene list enrichment analysis. **B.** Ten most downregulated gene pathways in PD *in vitro* differentiated microglia compared to PD transplanted microglia, as assessed by gene list enrichment analysis. n=3 cell lines per group, 2-3 transplanted brains per cell line or 3 biological experiments per cell line.

**Figure S4. Downregulated gene pathways in human microglia differentiated *in vitro* vs transplanted human microglia. A.** Ten most upregulated gene pathways in control *in vitro* differentiated microglia compared to control transplanted microglia, as assessed by gene list enrichment analysis. **B.** Ten most upregulated gene pathways in PD *in vitro* differentiated microglia compared to PD transplanted microglia, as assessed by gene list enrichment analysis. n=3 cell lines per group, 2-3 transplanted brains per cell line or 3 biological experiments per cell line.

**Figure S5. Transplanted PD striatal microglia display similar levels of activation compared to control in non-inflammatory conditions (no LPS challenge). A.** Separate percentages of ramified, bushy I, bushy II and ameboid cells in PD and control transplanted microglia in non-inflammatory conditions (no LPS injection) in the three control/PD isogenic pairs used in this study. Repeated-measures two-way ANOVA followed by Sidak's multiple comparisons test. N= 3-5 mice, 35 to 294 cells per mouse. **B.** Separate percentages of CD68+ cells in transplanted PD and control microglia in non-inflammatory conditions for the three control/PD isogenic pairs used in this study. Unpaired t-test. N= 3 mice, 36 to 179 cells per mouse. **C.** Integrated density of Iba1 signal in each isogenic control/PD pair in non-inflammatory conditions. Unpaired Mann-Whitney test. N= 12 to 100 cells per mouse, 3 mice per cell line. **D.** Separate percentages of p65-positive cells in transplanted PD and control microglia in non-inflammatory conditions for the three control/PD isogenic pairs used in this study. N= 3 mice, 72 to 215 cells per mouse. The PD 1 cell line displayed a decreased percentage of p65+ microglia compared to Control 1, however the difference was not observed in the two other pairs. Unpaired t-test. Data is represented as mean±SEM. *: $p<0.05$

**Figure S6. Transplanted PD striatal microglia displays increased pro-inflammatory activation compared to isogenic control in pro-inflammatory conditions. A.** Separate percentages of ramified, bushy I, bushy II and ameboid cells in PD and control transplanted microglia in pro-inflammatory conditions (after LPS challenge) in the three control/PD isogenic pairs used in this study. In all three pairs, a significant shift of microglia morphology towards more activated can be observed. Repeated-measures mixed-effects analysis followed by Sidak's multiple comparisons test. N= 3-4 mice, 74 to 239 cells per mouse. **B.** Separate percentages of CD68+ cells in transplanted PD and control microglia in pro-inflammatory conditions for the three control/PD isogenic pairs used in this study. In all three pairs, the percentage of CD68+ cells was significantly increased in PD microglia compared to control. Unpaired t-test. N= 3-4 mice, 24 to 239 cells per mouse. **C.** Integrated density of Iba1 signal in each isogenic control/PD pair after LPS challenge. In all three pairs, the integrated density of Iba1 signal was significantly increased in PD microglia compared to control. Unpaired Mann-Whitney test. N= 3-4 mice, 25 to 121 cells per mouse. **D.** Separate percentages of p65+ cells in transplanted PD and control microglia in pro-inflammatory conditions for the three control/PD isogenic pairs used in this study. In all three pairs, the percentage of p65+ cells was significantly increased in PD microglia compared to control. Unpaired t-test. N= 3-4 mice, 46 to 461 cells per mouse. Data is represented as mean±SEM. *: $p<0.05$; **: $p<0.01$; ***: $p<0.001$; ****: $p<0.0001$.

**Figure S7. Transplanted PD striatal microglia displays strongly decreased catalase expression and increased oxidative stress. A.** Separate integrated density measures of catalase signal in the three control/PD isogenic pairs used in this study in non-inflammatory conditions. In all three pairs, a strong decrease in the integrated density of catalase signal can be observed. Unpaired Mann-Whitney test. N= 12 to 100 cells per mouse, 3 mice per cell line. **B.** Separate integrated density measures of 8-hydroxy-2'-deoxyguanosine (8-OHdG) signal in the three control/PD isogenic pairs used in this study, in non-inflammatory conditions. In all three pairs, a strong increase in the integrated density of 8-OHdG signal can be observed. Unpaired Mann-Whitney tests. N=20 to 141 cells, 3-6 mice per cell line. **C.** Separate integrated density measures of 8-hydroxy-2'-deoxyguanosine (8-OHdG) signal in the three control/PD isogenic pairs used in this study, in pro-inflammatory conditions (LPS injection). In all three pairs, a strong increase in the integrated density of 8-OHdG signal can be observed. Unpaired Mann-Whitney tests. N=30 to 121 cells, 3-4 mice per cell line. **D.** Separate integrated density measures of catalase signal in the three control/PD isogenic pairs used in this study, in pro-inflammatory conditions (LPS injection). N=34 to 160 cells, 3-4 mice per cell line. Unpaired Mann-Whitney tests. Data is represented as mean±SEM. *: $p<0.05$; **: $p<0.01$; ***: $p<0.001$; ****: $p<0.0001$.

**Figure S8. Transplanted PD and control striatal microglia have low and similar levels of apoptosis. A.** Confocal maximum intensity projection of transplanted human microglia at 2 months post-transplantation, immunostained for cleaved caspase-3. Scale bar is 20 µm. **B.** Average percentage of cleaved caspase-3-positive cells in control and PD transplanted striatal microglia in non-inflammatory conditions (no LPS challenge). N=3 cell lines per group, 3 mice per cell line. **C.** Separate percentages of cleaved caspase-3-positive cells in control and PD transplanted microglia for the three isogenic pairs used in this study in non-inflammatory conditions. N=3 mice per cell line, 70 to 232 cells per mouse. **D.** Average percentage of cleaved caspase-3-positive cells in control and PD transplanted striatal microglia in pro-inflammatory conditions (LPS injection). N=3 cell lines per group, 3 mice per cell line. **E.** Separate percentages of cleaved caspase-3-positive cells in control and PD transplanted microglia for the three isogenic pairs used in this study in pro-inflammatory conditions (LPS injection). N=3 mice per cell line, 66 to 166 cells per mouse. Data is represented as mean±SEM. Unpaired t-tests.

**Figure S9. Transplanted PD and control striatal microglia have low and similar levels of cell division. A.** Confocal maximum intensity projection of transplanted striatal human microglia at 2 months post-transplantation, immunostained for ki67, a marker of cell proliferation. Scale bar is 20 µm. **B.** Average percentage of ki67-positive cells in control and PD transplanted striatal microglia in non-inflammatory conditions (no LPS injection). N=3 cell lines per group, 3 mice per cell line. **C.** Separate percentages of ki67-positive cells in control and PD transplanted striatal microglia for the three isogenic pairs used in this study in non-inflammatory conditions. N=3 mice per cell line, 61 to 275 cells per mouse. **D.** Average percentage of ki67-positive cells in control and PD transplanted striatal microglia in pro-inflammatory conditions (LPS injection). N= 3 cell lines per group, 3 transplanted mice per cell line. **E.** Separate percentages of ki67-positive cells in control and PD transplanted striatal microglia for the three isogenic pairs used in this study in pro-inflammatory conditions. N=3 mice per cell line, 24 to 239 cells per mouse. Data is represented as mean±SEM. Unpaired t-tests.

**Figure S10. Transplanted PD and control microglia have low and comparable levels of cellular senescence. A.** Confocal maximum intensity projection of transplanted human striatal microglia at 2 months post-transplantation, immunostained for p16, a marker of cellular senescence. Arrow shows a weakly stained p16-positive cell. Scale bar is 10 µm. **B.** Average percentage of p16-positive cells in control and PD transplanted striatal microglia in non-inflammatory conditions (no LPS injection). N=3 cell lines per group, 3 mice per cell line. **C.** Separate percentages of p16-positive cells in control and PD transplanted striatal microglia for the three isogenic pairs used in this study in non-inflammatory conditions. N=3 mice per cell line, 57 to 461 cells per mouse. **D.** Average percentage of p16-positive cells in control and PD transplanted striatal microglia in pro-inflammatory conditions (LPS injection). N=3 cell lines per group, 3 mice per cell line. **E.** Separate percentages of p16-positive cells in control and PD transplanted striatal microglia for the three isogenic pairs used in this study in pro-inflammatory conditions. Although the average percentage of p16-positive cells was increased in PD compared to control striatal microglia, because the separate percentages for the three pairs were not significant, we conclude that there is no significant difference between PD and control. N=3 mice per cell line, 46 to 476 cells per mouse. N= 3 cell lines per group, 3 transplanted mice per cell line. Data is represented as mean±SEM. Unpaired t-tests; *: $p<0.05$.

### Tables

**Table 1: Origin of the human pluripotent stem cell lines used in this study.** Cell line nomenclature is as in Soldner *et al.,* Cell 2011.

**Table 2: Primary antibodies**

**Table 3: Secondary antibodies**

**Table 4: All the cell lines have successfully differentiated into microglia at 29 days *in vitro*.** Percentages of Iba1-P2RY12-double positive cells per cell line. N= 3 independent experiments per cell line, 32 to 67 cells per experiment.

**Table 5: Ten most downregulated genes in *in vitro* differentiated PD microglia compared to isogenic control.** Gene ontologies of particular interest are in bold characters. N= 3 cell lines per group, 3 independent experiments per cell line.

**Table 6: Ten most upregulated genes in *in vitro* differentiated PD microglia compared to isogenic control.** Gene ontologies of particular interest are in bold characters. N= 3 cell lines per group, 3 independent experiments per cell line.

**Table 7: All the cell lines have successfully differentiated into microglia at 2 months after transplantation into the mouse brain.** Percentages of Iba1-P2RY12-double positive cells per cell line. N= 3 animals per cell line, 41 to 219 cells per animal.

**Table 8: Ten most downregulated genes in transplanted PD microglia compared to isogenic control.** Gene ontologies of particular interest are in bold characters. N= 3 cell lines per group, 3 independent experiments per cell line.

**Table 9: Ten most upregulated genes in transplanted PD microglia compared to isogenic control.** Gene ontologies of particular interest are in bold characters. N= 3 cell lines per group, 3 independent experiments per cell line.

### Supplemental tables

**Table S1**: Downregulated gene pathways in PD versus isogenic control *in vitro* differentiated microglia, using the Hallmark gene set collection from MSigDB24 (ftp.broadinstitute.org://pub/gsea/msigdb/human/gene_sets/h.all.v2023.1.Hs.symbols.gmt).

**Table S2**: Upregulated gene pathways in PD versus isogenic control *in vitro* differentiated microglia, using the Hallmark gene set collection from MSigDB24 (ftp.broadinstitute.org://pub/gsea/msigdb/human/gene_sets/h.all.v2023.1.Hs.symbols.gmt).

**Table S3**: Downregulated gene pathways in PD versus isogenic control *in vitro* differentiated microglia, using the curated gene set collection C2 from MSigDB24 (ftp.broadinstitute.org://pub/gsea/msigdb/human/gene_sets/c2.all.v2023.1.Hs.symbols.gmt).

**Table S4**: Upregulated gene pathways in PD versus isogenic control *in vitro* differentiated microglia, using the curated gene set collection C2 from MSigDB24 (ftp.broadinstitute.org://pub/gsea/msigdb/human/gene_sets/c2.all.v2023.1.Hs.symbols.gmt).

**Table S5**: Downregulated gene pathways in PD versus isogenic control transplanted microglia, using the Hallmark gene set collection from MSigDB24 (ftp.broadinstitute.org://pub/gsea/msigdb/human/gene_sets/h.all.v2023.1.Hs.symbols.gmt).

**Table S6**: Upregulated gene pathways in PD versus isogenic control transplanted microglia, using the Hallmark gene set collection from MSigDB24 (ftp.broadinstitute.org://pub/gsea/msigdb/human/gene_sets/h.all.v2023.1.Hs.symbols.gmt).

**Table S7:** Downregulated gene pathways in PD versus isogenic control transplanted microglia, using the curated gene set collection C2 from MSigDB24 (ftp.broadinstitute.org://pub/gsea/msigdb/human/gene_sets/c2.all.v2023.1.Hs.symbols.gmt)

**Table S8**: Upregulated gene pathways in PD versus isogenic control transplanted microglia, using the curated gene set collection C2 from MSigDB24 (ftp.broadinstitute.org://pub/gsea/msigdb/human/gene_sets/c2.all.v2023.1.Hs.symbols.gmt)
